# Transcriptional and phenotypic heterogeneity underpinning venetoclax resistance in AML

**DOI:** 10.1101/2024.01.27.577579

**Authors:** Vakul Mohanty, Natalia Baran, Yuefan Huang, Cassandra L Ramage, Laurie M Cooper, Shan He, Ramiz Iqbal, May Daher, CTD2 Research Network, Jeffrey W. Tyner, Gordon B. Mills, Marina Konopleva, Ken Chen

## Abstract

The venetoclax BCL2 inhibitor in combination with hypomethylating agents represents a cornerstone of induction therapy for older AML patients, unfit for intensive chemotherapy. Like other targeted therapies, venetoclax-based therapies suffer from innate and acquired resistance. While several mechanisms of resistance have been identified, the heterogeneity of resistance mechanism across patient populations is poorly understood. Here we utilized integrative analysis of transcriptomic and *ex-vivo* drug response data in AML patients to identify four transcriptionally distinct VEN resistant clusters (VR_C1-4), with distinct phenotypic, genetic and drug response patterns. VR_C1 was characterized by enrichment for differentiated monocytic- and cDC-like blasts, transcriptional activation of PI3K-AKT-mTOR signaling axis, and energy metabolism pathways. They showed sensitivity to mTOR and CDK inhibition. VR_C2 was enriched for *NRAS* mutations and associated with distinctive transcriptional suppression of *HOX* expression. VR_C3 was characterized by enrichment for *TP53* mutations and higher infiltration by cytotoxic T cells. This cluster showed transcriptional expression of erythroid markers, suggesting tumor cells mimicking erythroid differentiation, activation of JAK-STAT signaling, and sensitivity to JAK inhibition, which in a subset of cases synergized with venetoclax. VR_C4 shared transcriptional similarities with venetoclax-sensitive patients, with modest over-expression of interferon signaling. They were also characterized by high rates of *DNMT3A* mutations. Finally, we projected venetoclax-resistance states onto single cells profiled from a patient who relapsed under venetoclax therapy capturing multiple resistance states in the tumor and shifts in their abundance under venetoclax selection, suggesting that single tumors may consist of cells mimicking multiple VR_Cs contributing to intra-tumor heterogeneity. Taken together, our results provide a strategy to evaluate inter- and intra-tumor heterogeneity of venetoclax resistance mechanisms and provide insights into approaches to navigate further management of patients who failed therapy with BCL2 inhibitors.

## Introduction

AML (Acute Myeloid Leukemia) cells often upregulate the anti-apoptotic protein BCL2 to avoid apoptosis, thus targeting BCL2 represents an attractive therapeutic opportunity in AML^1^. Treating AML with a combination of venetoclax (VEN), a BCL2 inhibitor, and hypomethylating agents like azacitidine (AZA), is highly effective, especially in older patients who cannot tolerate intensive conventional therapy^2–4^. However, as is the case with other targeted therapies, VEN-based therapies are associated with frequent primary and acquired resistance^4,5^, with long-term cure rates of less than 25%. Considerable efforts have been made to identify mechanisms of resistance in VEN-based therapies and strategies to overcome them^1,5^. The efficacy of VEN therapies has been linked to their ability to eliminate leukemia stem cells (LSCs), that have capacity for self-renewal^6^. AML LSCs are dependent on OxPhos^6^ driven by upregulating amino acid (AA) metabolism^7^. Combination of VEN and AZA suppresses AA uptake^7^, resulting in cell death. Consequently, activation of alternative energy metabolism pathways that can compensate for VEN mediated suppression of OxPhos result in resistance and present a viable target to eradicate VEN resistant LSCs^8,9^. CRISPR (Clustered Regularly Interspaced Short Palindromic Repeats) screening in a lymphoma cell line has also linked increase in OxPhos through dysregulation of AMPK signaling to VEN resistance^10^. CRISPR screens have also linked mitochondrial structure and function to VEN resistance^11,12^. Disruption of mitochondrial structure^11^ and translation^12^ result in induction of stress response pathways that re-sensitizes cells to VEN. Intriguingly loss of TP53 expression, which has been linked to VEN resistance is also accompanied by increased OxPhos^13^.

Recent studies have linked differentiation status of AML cells to VEN resistance. Using clinical samples, Pei et al.^14^ showed that monocytic AML (more differentiated monocytic(Mono)-like AML cells) suppress BCL2, have high expression of OxPhos genes and rely on MCL1 for energy metabolism and survival, rendering them inherently resistant to VEN+AZA therapy. Emergence of monocytic clones can occur upon relapse^14^. Correlating deconvoluted abundances of blasts, delineated by development status, from bulk-RNAseq with *ex-vivo* drug responses showed association of VEN resistance with enrichment of Mono-like blasts^15,16^. Enrichment of monocytic signature was also observed in primary human AML specimens resistance to VEN *ex vivo*^17^. However, data from a clinical trial^18^ did not recapture this association and recent study by Waclawiczek et al.^19^ suggests that resistance to VEN+AZA is predictable from expression patterns of BCL2 family of proteins in LSCs rather than presence of Mono-like sub-populations. A recent study has also linked VEN resistance to erythroid (FAB(French-American_British classification)-M6) and megakaryocytic (FAB-M7) leukemias, driven by over-expression of BCL2L1^20^. Taken together, this body of work suggests that developmental characteristics of AML blasts can influence sensitivity to VEN-based therapies. As described above differentiated AML blasts often over-express other anti-apoptotic proteins like MCL1, BCL2L1 and BCL2A1 that can mitigate efficacy of VEN-based therapies^1,5,21^. While their inhibition presents an approach to overcome VEN resistance *in vitro*^20,21^, toxicities associated with their inhibition has limited their clinical application^5,22,23^. Thus, there is a need to identify alternative approaches to target VEN resistant differentiated blasts.

While significant efforts in the field have revealed various mechanisms of resistance to VEN therapy, heterogeneity of these mechanisms in patient population and associated therapeutic vulnerabilities are poorly understood. Analysis of patient cohorts have linked common mutations in AML such as tandem duplication in *FLT3* (*FLT3-ITD*), *TP53*, *RAS* and *PTPTN1* among others^21,24,25^ to VEN resistance. Indicating diverse contribution of genetic alterations to VEN resistance, which might point to associated mechanistic heterogeneity. However, integrative multi-omics analysis has not been performed across large-scale AML cohorts to elucidate VEN resistance states. In this study, we used integrative analysis of bulk-RNAseq and *ex-vivo* drug response data to identify four transcriptionally distinct clusters/states of VEN resistant patients. These states have distinct mutational, phenotypic and drug response characteristics, capturing inter-tumor heterogeneity associated with VEN resistance. Using single cell RNAseq (scRNAseq), we also illustrate that multiple resistance states can be present in a single tumor and can therefore facilitate interrogation of intra-tumor heterogeneity associated with VEN resistance.

## Results

### Decomposing VEN resistance gene-expression signature in VEN resistant patients to capture inter-patient heterogeneity

To explore transcriptional heterogeneity underlying VEN resistance in AML, we first derived a gene-expression signature associated with VEN resistance from cell-lines (**Figure 1A**), with homogeneously defined relationships between gene expressions and drug responses. Briefly, a Gaussian mixture model was fitted to VEN AUC (Area Under Curve) values of leukemia cell lines to identify VEN sensitive and resistant cell lines (**Figure 1A left** and **S1B,** see **Methods**). Differential expression analysis comparing resistant and sensitive cell-lines identified 1297 differentially expressed genes (DEGs, q < 0.1 and absolute log2 fold-change (log2FC) > 1; **Figure 1A right**). The signature was enriched for genes in Heme metabolism, *KRAS* signaling (down) and myogenesis pathways (**Figure S1C**, q< 0.1). Genes in the signature with low variance and mean expression in patient samples from BeatAML1^26^ were filtered out (see **Methods**). Post-filtering (613 genes), the signature retained its functional enrichment profile (**Figure S1C**). Next, VEN resistant patients (VRPs) in BeatAML1 were identified (see **Methods**). We extracted normalized expression levels of the signature genes and decomposed them into constituent factors and their contribution in each sample using non-negative matrix factorization (NMF) (**Figure 1B** and see **Methods**). The NMF components described different gene expression programs, constituting the resistance signature, and loadings of the components reflecting their respective contribution in each sample. The patient loadings in these programs were scaled and clustered, using consensus clustering, to identify four clusters of VRPs (VEN resistant clusters (VR_Cs) 1-4; **Figure 1C**, **Table 1**, see **Methods**). We refer to the VR_Cs along with the VEN sensitive patients, or VSC (VEN sensitive cluster), collectively as the five VEN responsive states (VRS) in the following sections.

**Figure 1:**
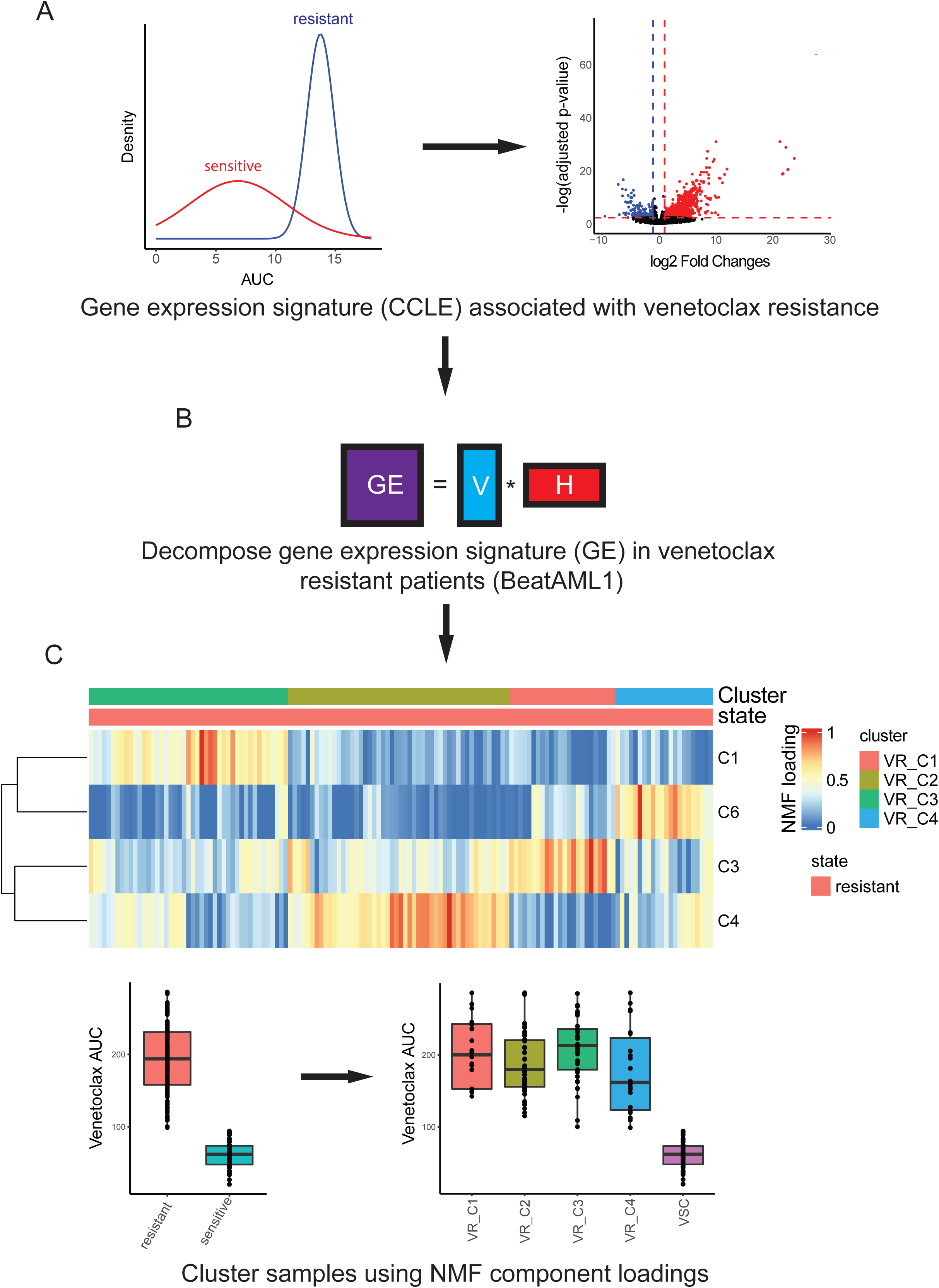
Workflow to identify transcriptionally distinct VEN resistant patients: **A)** A Gaussian mixture model is fit to VEN AUC of cell-lines to identify sensitive and resistant cell-lines (left; see **Methods**). Resistant and sensitive cell-lines are compared to identify differentially expressed genes (right). This gene expression signature (GE) is used for decomposition in **B**. **B)** NMF decomposition of GE in VEN resistant samples in BeatAML1 **V** and **H** are gene and patient loading matrices of NMF components **C)** top: VR_Cs identified by clustering **H**, (see **Methods)**. bottom: boxplots of VEN AUC in resistant and sensitive patients (left) and across VRS (right).

**Table 1:**
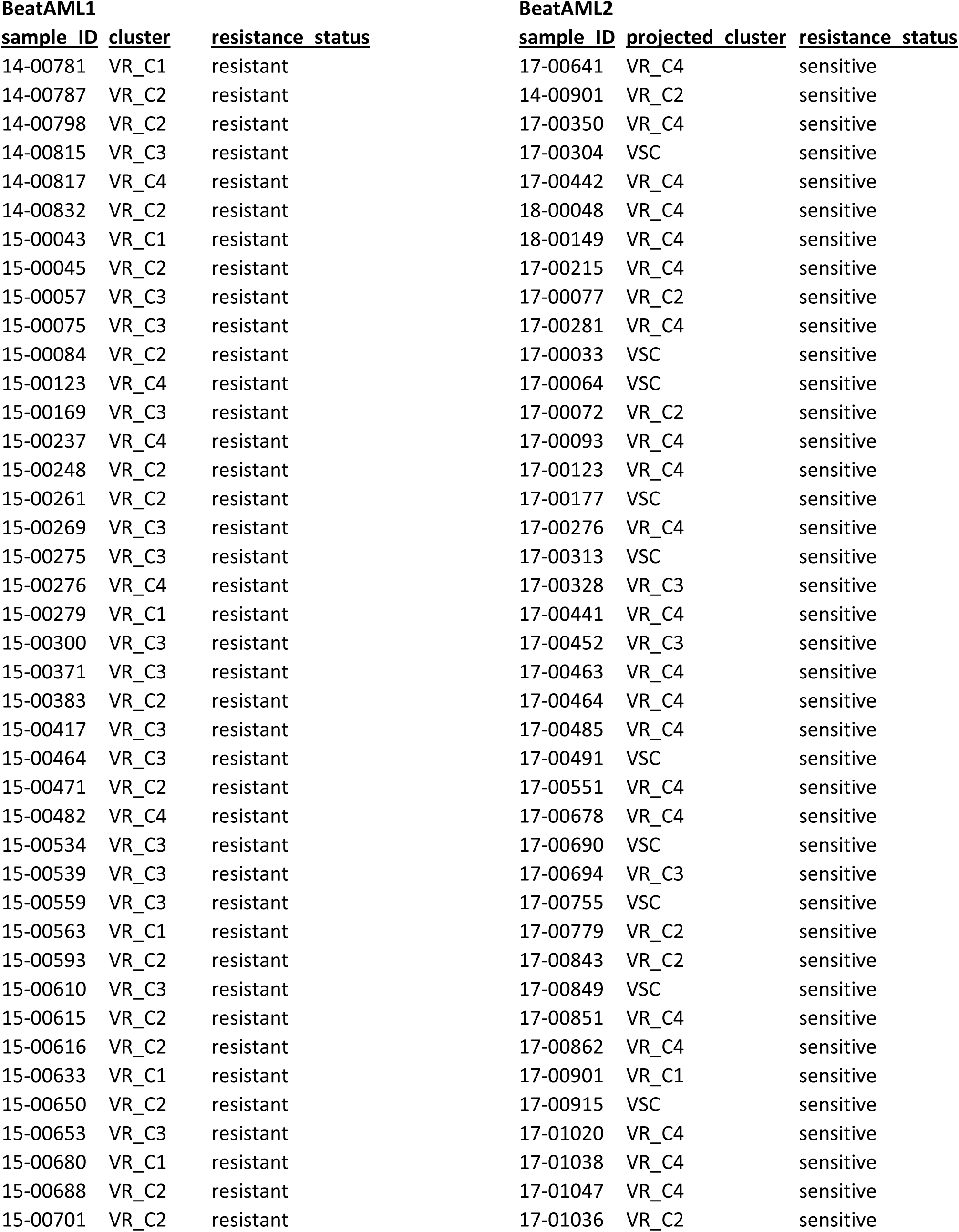
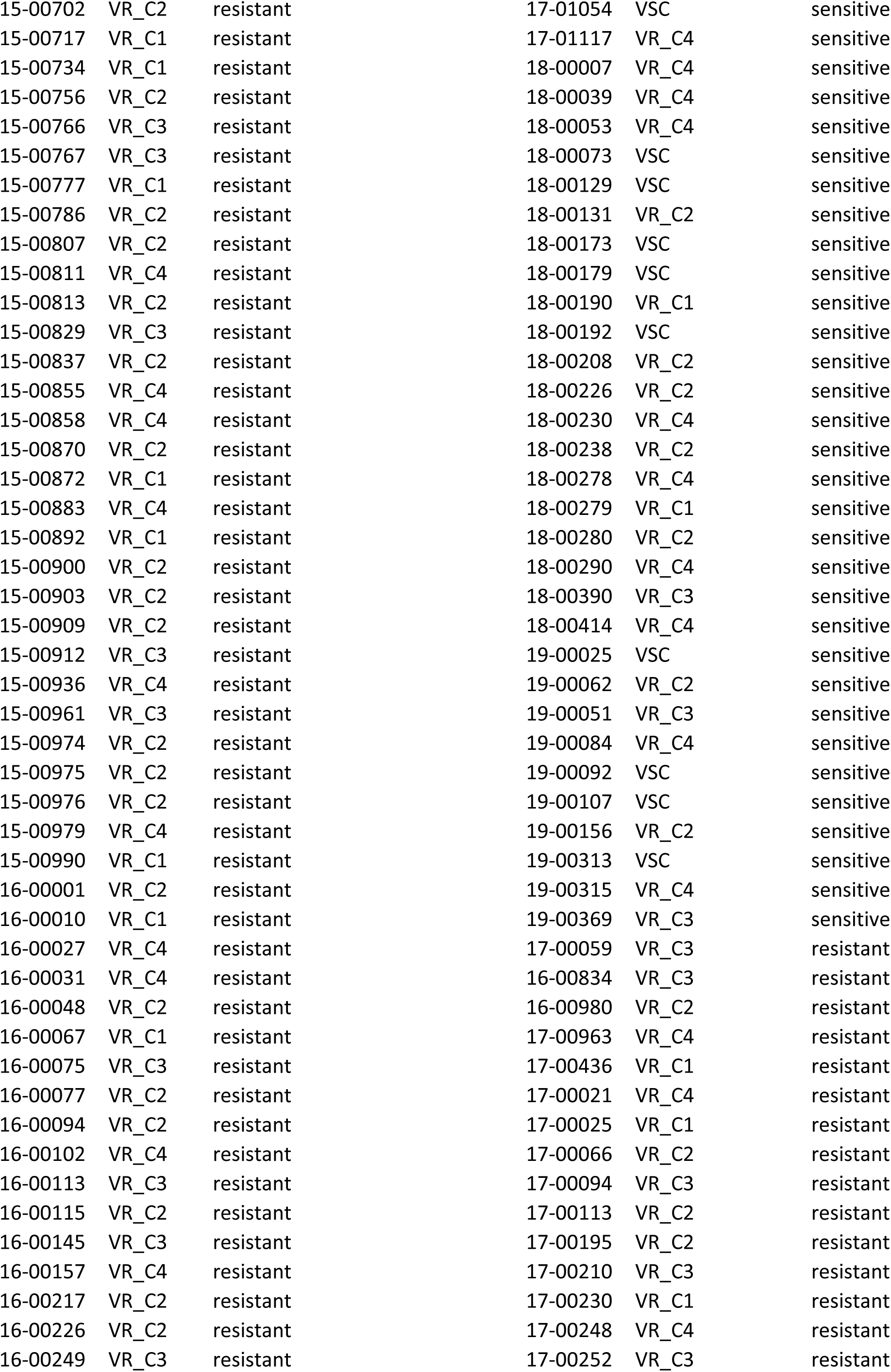
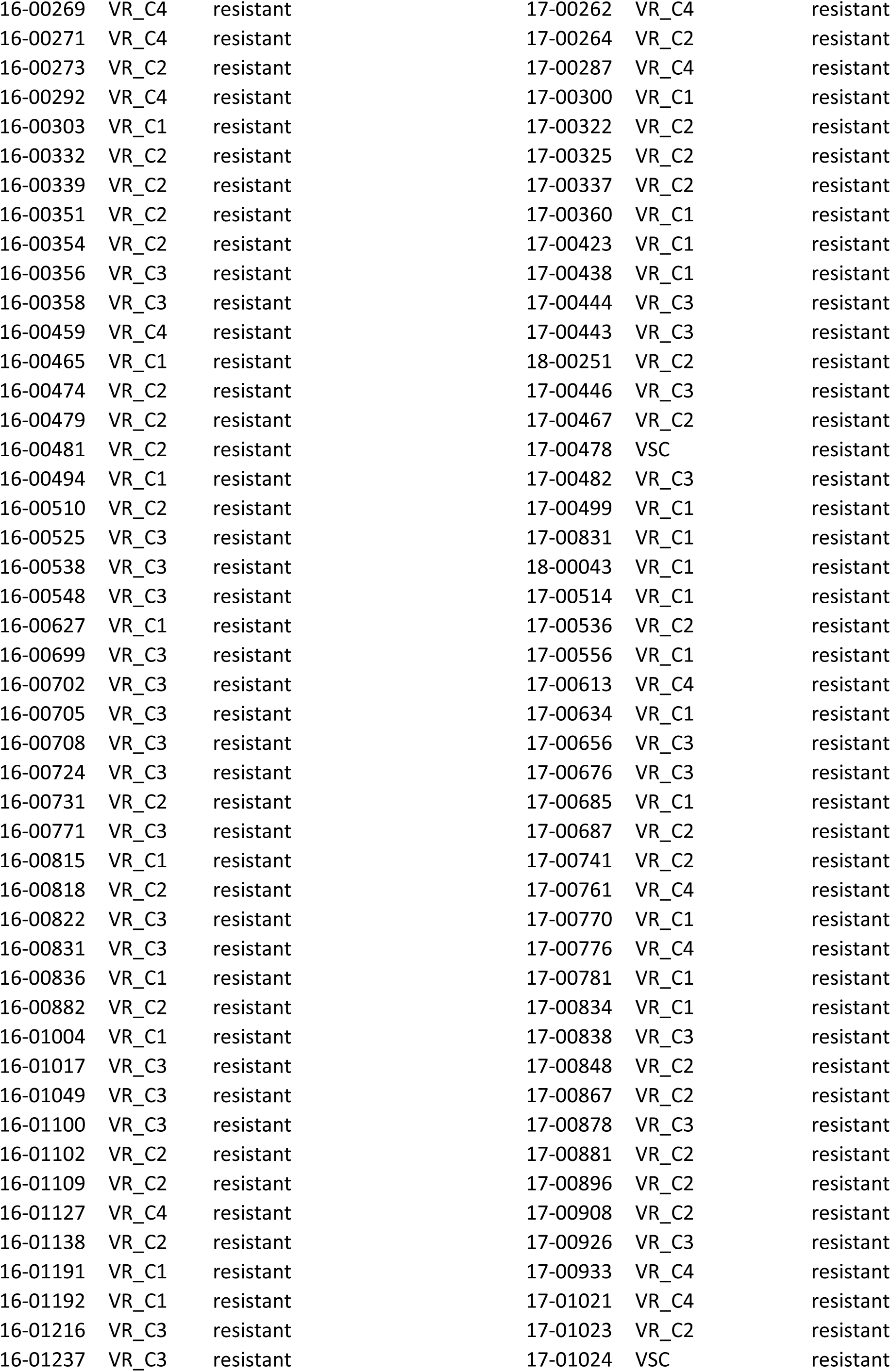
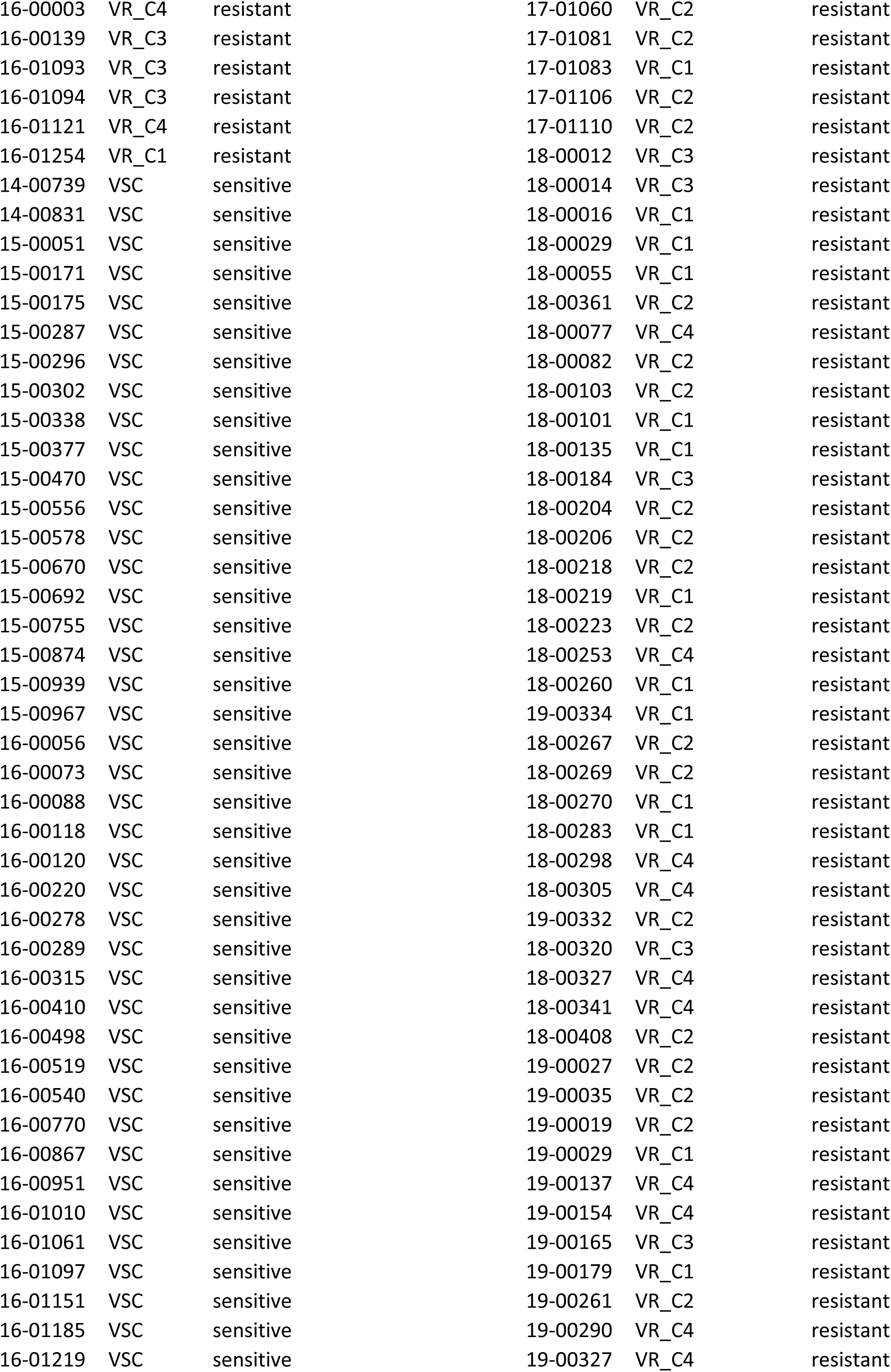
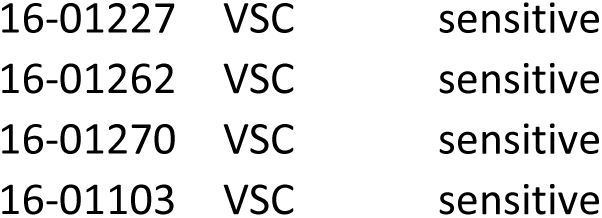
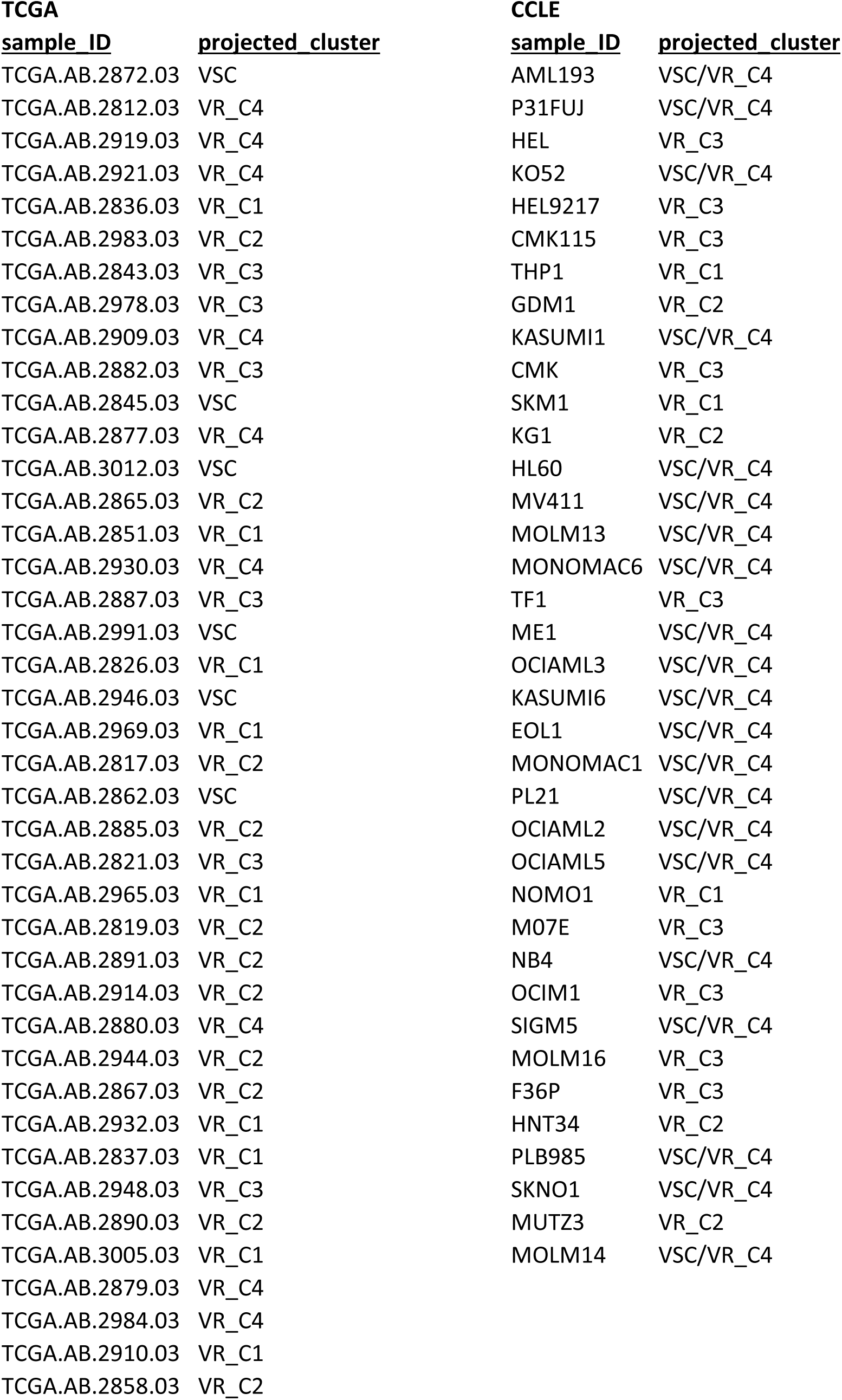
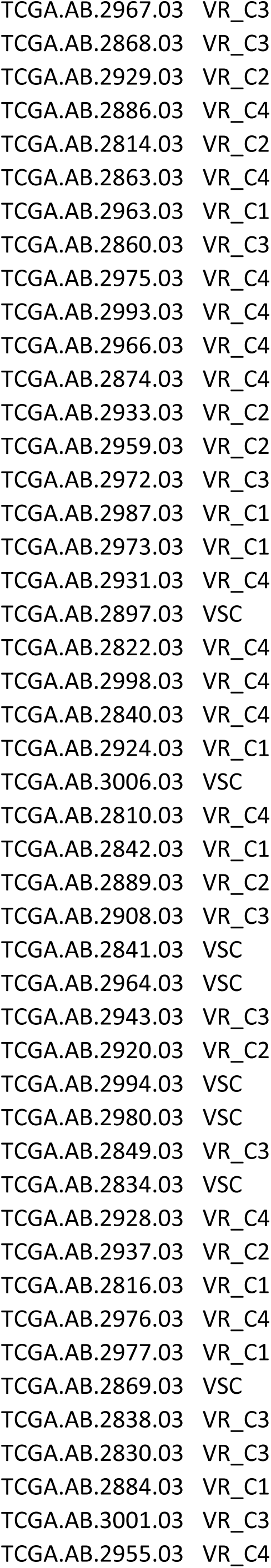
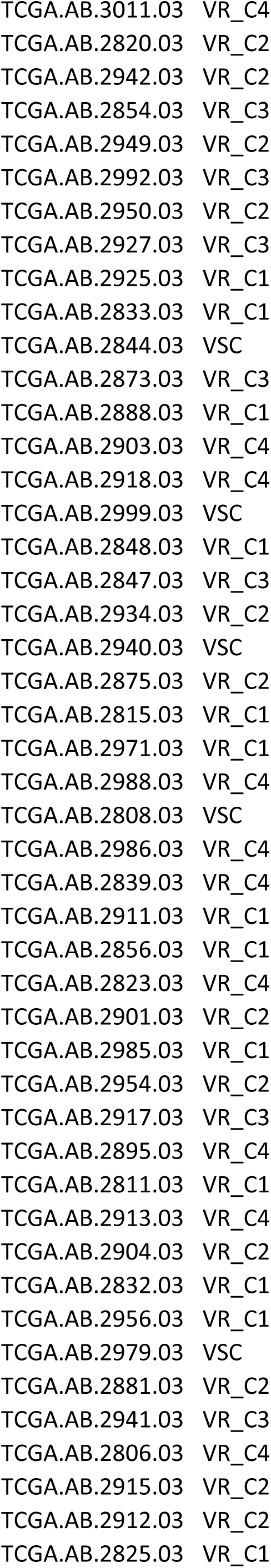
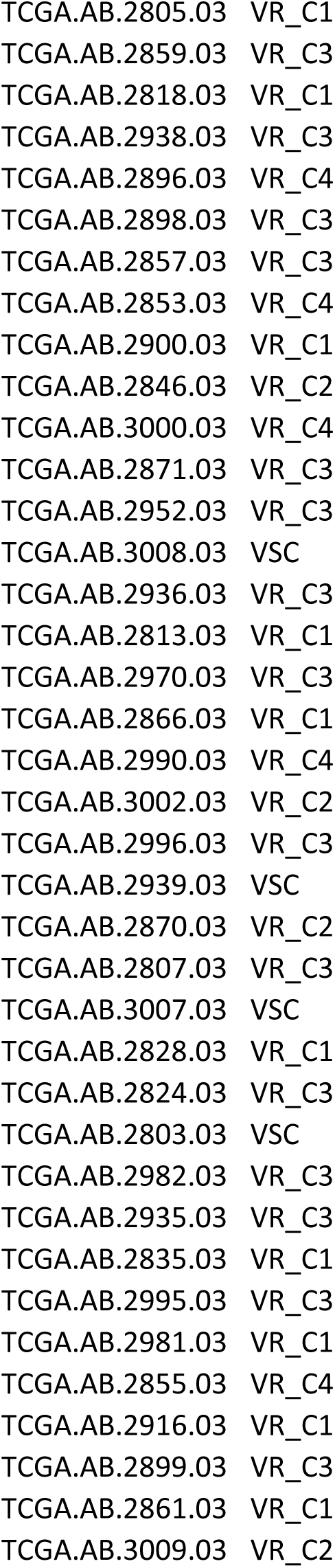
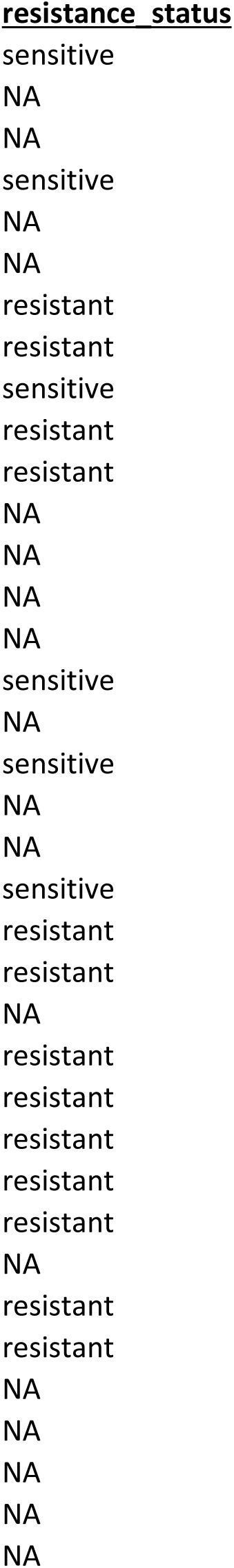
Cluster membership for original clustering in BeatAML1 and projected clustering (BeatAML2, TCGA.

### Clinical and genetic characteristics of VRS

We compared clinical characteristics between VRS to assess phenotypic differences (**Figure 2A-B**). We identified several clinical factors that show variability across VRS (q < 0.1; **Figure 2A**). VR_C1 was characterized by the high monocytic % in peripheral blood (PB) (**Figure 2A**) and enrichment for FAB_M5 patients (72%, **Figure 2C**) indicating enrichment for Mono-like blasts. VR_C1 also showed elevated levels of lactate dehydrogenase (LDH), aspartate transaminase (AST) and mean corpuscular volume (MCV) (**Figure 2A**), consistent with tissue damage, liver, and bone marrow (BM) dysfunction, respectively. VR_C3, in contrast, was characterized by patients who were older at diagnosis, had low PB blast %, low white blood cell (wbc) counts, with high neutrophil % and lymphocytes % in PB (**Figure 2A**). We also observed modest enrichment (∼36%) for residual samples (i.e., cancer remaining after chemotherapy). VR_Cs also showed specific and prominent risk distinctions as defined by European LeukemiaNet (ELN)2017 classification: with VSC and VR_C4 showing enrichment for patients with favorable classification (∼ 36% and 41% respectively), VR_C3 for patients classified as non-initial (i.e., specimen is not at initial diagnosis when ELN was accessed; 51%), and VR_C2 showed a modestly higher rate of patients classified with adverse risk (30%) and FAB_M2 (50%, **Figure 2C**). Further, we also observed distinct patterns of clinical characteristics between VR_Cs and VSC (adjusted p-value < 0.1; **Figure 2B**): All VR_Cs except for VR_C4 showed significantly lower blast % in PB and BM relative to VSC, while other clinical variables like monocytic % in PB, age at diagnosis and lymphocyte % in PB showed cluster specific differential patterns (**Figure 2B**). While correlation of clinical variables to VEN AUC have been previously reported^21^, our analysis localized these associations to specific patient clusters.

**Figure 2:**
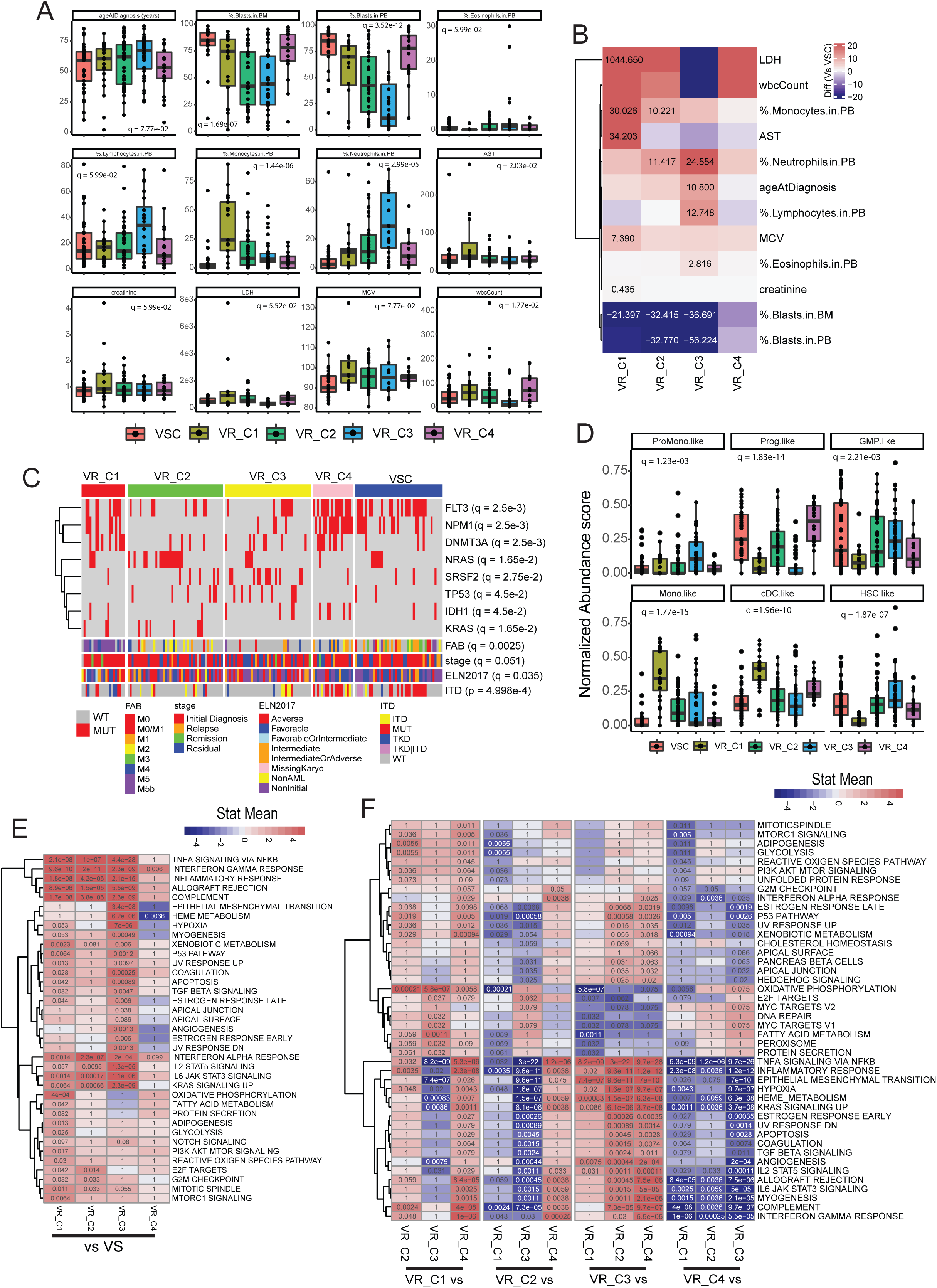
Phenotypic, genetic, and transcriptional characteristics of VRS (in BeatAML1): **A)** Boxplot of clinical variables significantly different (q < 0.1) between VRS. **B)** Heatmap of difference between means of clinical variables (in **A**) in each VR_C (column) relative to VSC. The difference is reported for significant comparisons (adjusted P-values (p.adj) < 0.1) **C)** Heatmap of mutations and clinical factors showing significant association with VRS (q < 0.1, see **Methods**). **D)** Abundance scores of AML blast**-**types with significantly different (q < 0.1) abundances across VRS. Heatmap of differentially expressed pathways comparing VR_Cs to VSC **(E)** and each other **(F)**. Red indicates over-expression and blue suppression. Text in each cell is the q-value, with significance at q < 0.1. **Note:** comparison of differences between multiple groups was performed using ANOVA and p-values corrected by FDR. Post-hoc pairwise testing was performed using Tukey’s test.

VRS also showed distinct mutational patterns (q < 0.1; **Figure 2C**); *FLT3* and *NPM1* are more frequently mutated in VSC, VR_C1 and 4. VR_C1 and 4 were further enriched for mutations in *DNMT3A*, while *IDH1* mutations were found mainly in VR_C4 (**Figure 2C**). VR_C1 also showed a strong preference for *FLT3*-TKD mutations (4/9 mutants). Of the FTL3-ITD mutants, all but one was spread across resistant clusters (VR_C2-4). RAS mutations, which have been associated with VEN resistance^14,21^ were frequently present in VR_C2 (*NRAS* (∼35%) and *KRAS* (∼14%)) and showed modest mutation rates in VR_C1 (∼14% for both). *NRAS* mutations in VR_C2 were also mutually exclusive from *FTL3* mutations (**Figure S2A**, top). VR_C3 was characterized by mutations in *DNMT3A (*16%), *FLT3* (19%), *TP53* (19%) and *SRSF2* (28%) (**Figure 2C**). Of note, *TP53* mutants in VR_C3 were mutually exclusive from mutations in *SRSF2* and *FLT3* (**Figure S2A**, bottom).

These data indicate that the VR_Cs derived from transcriptomic data have distinct phenotypic and genomic characteristics, further underlining the heterogeneity associated with VEN resistance.

### Cell type composition of VRS

VEN resistance has been linked to differentiation status of blasts^14–16^. In agreement with these findings, sensitivity to VEN was positively correlated with the abundance of less differentiated progenitor-like (Prog-like) blasts and inversely with abundance of more differentiated Mono-like blasts and cDC-like blasts (q< 0.1; **Figure S2B**). Resistant samples also showed enrichment of differentiated blasts and depletion of prog-like blasts (q < 0.1; **Figure S2C**). However, cell-type composition varied among VR_Cs (**Figure S2D**). While there was a general trend for enrichment for Mono-like blasts in VR_Cs relative to VSC, the difference was most prominent in VR_C1, along with enrichment for cDC-like blasts (**Figure 2D** and **Figure S2E**), which agreed with the enrichment of FAB-M5 tumors in VR_C1 (**Figure 2C**). The depletion in prog-like blasts relative to VSC was restricted to VR_C1 and 3 (**Figure 2D** and **S2E**). Interestingly, VR_C2 and 3 had significantly higher fractions of less differentiated hematopoietic stem cell (HSC) and granulocyte-monocyte progenitor (GMP)-like blasts compared to VR_C1 with levels being comparable to VSC (**Figure 2D** and **S2E**). VR_C4 was characterized by high prevalence of progenitor-like blasts, like VSC, along with high cDC-like blasts (**Figure 2D** and **S2E**).

These data suggest that while we recapture the association of VEN resistance with high prevalence of monocytic blasts, the cell-type composition of VR_Cs varies. VR_C1 strongly recapitulates previously described^15,16^ enrichments for monocytic blasts and depletion of stem cell-like blasts in VEN resistant patients (**Figure S2E**). In contrast, other VR_Cs showed relative enrichment for more immature cell types as in VR_C4 (Prog-like) and VR_C3 (HSC and GMP-like; **Figure S2E**).

### Transcriptional characteristics of VRS

We next accessed differential activity of pathways in the VR_Cs using differential expression and gene-set enrichment analysis (GSEA)^27^. VR_Cs as a collective were characterized by increased activity of inflammatory pathways (Interferon signaling, interleukin(IL)2-signal transducer and activator of transcription (STAT)5 signaling, IL6-STAT3 signaling, tumor necrosis factor (TNF)α signaling, allograft rejection etc.), modest activation of cell surface components/signaling (apical surface, apical junction and notch signaling) and other pathways like apoptosis, kirsten rat sarcoma viral oncogene homolog (KRAS) signaling, TP53 pathway, epithelial mesenchymal transition (EMT), angiogenesis and hypoxia associated with malignant transformation and progression relative to VSC (**Figure S2F**). When the comparison was broken down to level of individual VR_Cs, many of the inflammatory pathways (IL2-STAT5 signaling, IL6-STAT3 signaling, TNFα signaling, allograft rejection, interferon signaling) and KRAS signaling up were upregulated across VR_Cs (**Figure 2E**). Interestingly, VR_C4 that had the lowest VEN AUC of the VR_Cs showed few differences relative to VSC unlike the other VR_Cs, which indicates a degree of transcriptional similarity between VR_C4 and VSC.

While these results recapitulated many of the general trends of pathway activity in VR_Cs as a whole (**Figure S2F**), we also identified cluster specific difference between individual VR_Cs and VSC (**Figure 2E**), as well as differences between VR_Cs (**Figure 2F**). VR_C1 showed specific activation of several metabolic pathways (glycolysis, OxPhos and fatty acid metabolism (FAM)) and activation of the (phosphatidylinositol 3-kinase) PI3K_mTOR pathways (PI3K-AKT-MTOR and MTORC1 signaling), which is a regulator of metabolic pathways^28^, relative to VSC (**Figure 2E**). These trends were also captured relative to other VR_Cs, with VR_C1 showing high OxPhos, FAM (vs VR_C2-3), glycolysis (vs VR_C2 and 4), PI3K-AKT-MTOR (vs VR_C2 and 4) and MTORC1 signaling (vs VR_C2 and 4) (**Figure 2F**). Comparing activity of signaling pathways inferred using *progeny*^29^ (see **Methods**), VR_C1 was characterized by activation of PI3K signaling relative to other VRS (q < 0.1, **Figure S2G-H**), consistent with GSEA.

VR_C2 did not show any unique pathway differentially regulated relative to VSC and other VR_Cs (**Figure 2E-F** and **S2H**). However, we observed suppression of transcriptional activity and expression of homeobox A cluster (*HOXA)* and homeobox B cluster (*HOXB)* genes (**Figure S2I-J**). While the mechanistic implications of this observation are unclear, suppression of HOX expression has previously been reported as a likely biomarker for resistance to VEN+AZA therapy^30^.

VR_C3 was characterized by activation of estrogen signaling, cell surface components (apical surface and apical junction) and EMT relative to VSC (**Figure 2E**), which could indicate perturbation in cell-cell signaling in the tumor microenvironment (TME). Many of these pathways were also over-expressed in VR_C3 relative to other VR_Cs (**Figure 2F**). VR_C3 furthermore showed a general trend of TP53 pathway activation, UV response and apoptotic signaling (**Figure 2E-F**). Dysregulation of these pathways in VR_C3 together with observed higher rate of TP53 mutation (**Figure 1C**) could indicate dysregulation of TP53 signaling and its regulation of DNA damage and apoptosis. Intriguingly, mutations^25^ and silencing^13^ of TP53 have been linked to VEN resistance. VR_C3 also showed activation of angiogenesis and hypoxia pathways relative to other clusters (**Figure 2F-G**), which was consistent with high hypoxia and (vascular endothelial growth factor) VEGF signaling inferred by *Progeny* (q < 0.1, **Figure S2G-H**). *Progeny* inferred signaling activity also indicated activation of several immune associated signaling pathways (JAK-STAT, TNFα, (Nuclear factor-κB) NFκB and TGF(transforming growth factor)β; q < 0.1, **Figure S2G-H**), consistent with the general trend of higher activation of hallmark immune pathways in VR_C3 relative to other VR_Cs (**Figure 2E-F**). VR_C3 was also characterized by higher infiltration of cytotoxic T-lymphocytes (CTL) and higher expression of cytotoxic effector and immune-checkpoint genes (**Figure S2L-K**). While VR_C1 also showed over-expression of immune pathways (**Figure 2E-F**), it did not show the infiltrated CTL phenotype (**Figure S2L-K**). Recently, it was reported that TP53 mutant AMLs have higher immune infiltration with high expression of checkpoint genes, making them responsive to immunotherapy^31^. Although enriched for TP53 mutants (19%, **Figure 2C**) compared to other VR_Cs, VR_C3 is predominantly TP53 WT (wild type), indicating other factors in addition to TP53 mutations influence induction of inflammatory signaling and immune cell infiltration.

VR_C4 showed few differences relative to VSC, with modestly high activation of Interferon signaling (**Figure 2E**) and no significant differences in activity of other signaling pathways (**Figure S2H**). Relative to other VR_Cs, its pathways profiles (**Figure 2F**) were reminiscent of those observed relative to VSC (**Figure 2E**). These indicate transcriptional similarities between VR_C4 and VSC. However, we did observe higher activity of OxPhos and MYC targets relative to VR_C3 (**Figure 2F**), which might indicate higher metabolic activity.

### VR_C clusters show distinct sensitivity to individual drugs

VR_Cs as a collective showed greater sensitivity to several drugs relative to VSC (see **Methods**, **Figure S2M and Table 2**). We next identified drugs that showed cluster specific sensitivity in each VR_C ((see **Methods**, **Figure S2N and Table 2**), recapturing many of the drugs with efficacy in the VR_Cs (**Figure S2M**). VR_C1 showed sensitivity to several PI3K-AKT-mTOR (BEZ235, rapamycin and GDC-0941) and RAF(rapidly accelerated fibrosarcoma kinase)_MEK(Mitogen-activated protein kinase/ERK kinase)_ERK(extracellular-signal-regulated kinase) inhibitors (trametinib and selumetinib). VR_C1 also showed sensitivity to inhibition of Hsp(heat shock protein)90 (tanespimycin), (histone deacetylase) HDAC (panabinostat), (cyclin-dependent kinase) CDK (SNS-032) and BRD(bromodomain protein)4 (JQ1). While other clusters shared sensitivity to inhibition of some of these pathways and proteins (PI3K-AKT-mTOR (VR_C4), RAF_MEK_ERK (VR_C2), CDK (VR_C3 and VR_C4) and Hsp90 (VR_C4)), they also showed distinct sensitivity to other inhibitors. VR_C3 and VR_C2 were selectively sensitive to Elesclomol (induces oxidative stress), while VR_C4 showed sensitivity to several tyrosine kinase inhibitors. Many of the pathways and proteins targeted by these drugs have been identified as potential targets to mitigate VEN resistance^5^. Our analysis suggests that many of these drugs are likely to be effective in specific AML subsets defined by VR_Cs (**Figure S2N**).

**Table 2:** Drugs that show specific sensitivity in VRPs, VR_C1 -4 in beatAML 1. q-values for h.

**BeatAML1**
| <u>drug</u> | <u>enrichment q</u> | <u>T-test q</u> | <u>cluster</u> |
| --- | --- | --- | --- |
| 17-AAG (Tanespimycin) | 0.023938197 | 0.008336 | VRPs |
| BEZ235 | 0.016193846 | 0.021796 | VRPs |
| Bortezomib (Velcade) | 0.010586678 | 0.004425 | VRPs |
| Dasatinib | 0.004978496 | 5.17E-05 | VRPs |
| Flavopiridol | 0.02678996 | 0.011314 | VRPs |
| GDC-0941 | 0.015982961 | 0.007526 | VRPs |
| INK-128 | 0.010713154 | 0.007555 | VRPs |
| JNJ-28312141 | 0.027752794 | 0.062181 | VRPs |
| KI20227 | 0.00477837 | 0.000372 | VRPs |
| PD173955 | 0.09201292 | 0.152286 | VRPs |
| PI-103 | 0.005528917 | 0.021796 | VRPs |
| PP242 | 0.016193846 | 0.26134 | VRPs |
| Panobinostat | 0.000124277 | 0.000372 | VRPs |
| Rapamycin | 0.005528917 | 4.61E-05 | VRPs |
| SNS-032 (BMS-387032) | 0.049654852 | 0.050023 | VRPs |
| Selumetinib (AZD6244) | 0.010586678 | 5.41E-06 | VRPs |
| Trametinib (GSK1120212) | 0.010586678 | 0.004636 | VRPs |
| AT7519 | 0.14664863 | 0.068772 | VRPs |
| Doramapimod (BIRB 796) | 0.111936211 | 0.050023 | VRPs |
| Elesclomol | 0.111936211 | 0.008953 | VRPs |
| JAK Inhibitor I | 0.545562751 | 0.081601 | VRPs |
| JQ1 | 0.111936211 | 0.021796 | VRPs |
| 17-AAG (Tanespimycin) | 0.127982184 | 0.017341 | VR_C1 |
| GDC-0941 | 0.003874683 | 0.035608 | VR_C1 |
| JQ1 | 0.188104085 | 0.071972 | VR_C1 |
| Panobinostat | 0.015461869 | 0.003385 | VR_C1 |
| SNS-032 (BMS-387032) | 0.130351637 | 0.024528 | VR_C1 |
| Selumetinib (AZD6244) | 0.127982184 | 0.003385 | VR_C1 |
| Trametinib (GSK1120212) | 0.025549786 | 0.064412 | VR_C1 |
| BEZ235 | 0.025549786 | 0.324063 | VR_C1 |
| Bortezomib (Velcade) | 0.080556956 | 0.247418 | VR_C1 |
| Rapamycin | 0.007612853 | 0.21272 | VR_C1 |
| Elesclomol | 0 | 3.74E-05 | VR_C2 |
| JQ11 | 0.049980164 | 0.042923 | VR_C2 |
| Selumetinib (AZD6244) | 0.224930667 | 0.071217 | VR_C2 |
| Trametinib (GSK1120212) | 0.004321798 | 0.071217 | VR_C2 |
| Elesclomol1 | 0.195405403 | 0.009712 | VR_C3 |
| SNS-032 (BMS-387032) | 0.195405403 | 0.038991 | VR_C3 |
| 17-AAG (Tanespimycin) | 0.005793033 | 0.000884 | VR_C4 |
| A-674563 | 0.007973172 | 0.015082 | VR_C4 |
| AT7519 | 0.128532888 | 0.051227 | VR_C4 |
| BEZ2351 | 0.510011351 | 0.082217 | VR_C4 |
| Dasatinib | 0.011306871 | 0.003601 | VR_C4 |
| Doramapimod (BIRB 796) | 0.04577057 | 0.030086 | VR_C4 |
| Dovitinib (CHIR-258) | 0.009161492 | 0.000884 | VR_C4 |
| Foretinib (XL880) | 0.000963125 | 8.19E-05 | VR_C4 |
| Gilteritinib (ASP-2215) | 0.562167546 | 0.054318 | VR_C4 |
| INK-128 | 0.032340234 | 0.002985 | VR_C4 |
| JAK Inhibitor I | 0.029891586 | 0.000153 | VR_C4 |
| JNJ-28312141 | 0.042232507 | 0.022633 | VR_C4 |
| KI20227 | 0.033440521 | 0.006044 | VR_C4 |
| PD173955 | 0.154265794 | 0.058263 | VR_C4 |
| PI-103 | 0.017708043 | 0.002532 | VR_C4 |
| PP242 | 0.08036522 | 0.072149 | VR_C4 |
| Pelitinib (EKB-569) | 0.00722152 | 0.003601 | VR_C4 |
| Ponatinib (AP24534) | 0.005793033 | 0.014875 | VR_C4 |
| Quizartinib (AC220) | 0.009161492 | 0.000884 | VR_C4 |
| Rapamycin | 0.238823927 | 0.031398 | VR_C4 |
| Tivozanib (AV-951) | 0.001802386 | 0.006698 | VR_C4 |

BeatAML2
| <u>drug</u> | <u>enrichment_q</u> | <u>T-test_q</u> | <u>cluster</u> |
| --- | --- | --- | --- |
| BEZ235 | 0.002886533 | 0.010313501 | VR_C1 |
| Bortezomib (Velcade) | 0.168509656 | 0.011026089 | VR_C1 |
| GDC-0941 | 0.001640075 | 0.015208708 | VR_C1 |
| Panobinostat | 1.34E-05 | 3.86E-11 | VR_C1 |
| Rapamycin | 9.47E-07 | 8.86E-06 | VR_C1 |
| Trametinib (GSK1120212) | 1.98E-05 | 0.000149204 | VR_C1 |
| 17-AAG (Tanespimycin) | 7.25E-05 | 1.93E-07 | VR_C1 |
| JQ1 | 0.168509656 | 0.620814291 | VR_C1 |
| SNS-032 (BMS-387032) | 0 | 0.006168604 | VR_C1 |
| Selumetinib (AZD6244) | 4.14E-05 | 8.68E-07 | VR_C1 |
| Elesclomol | 0 | 3.66E-08 | VR_C2 |
| JQ11 | 0.294075213 | 0.239357815 | VR_C2 |
| Trametinib (GSK1120212) | 0.02090248 | 0.060328228 | VR_C2 |
| Selumetinib (AZD6244) | 0.455290719 | 0.147067643 | VR_C2 |
| Elesclomol1 | 0.985271037 | 0.899919232 | VR_C3 |
| SNS-032 (BMS-387032) | 0.985271037 | 0.899919232 | VR_C3 |
| 17-AAG (Tanespimycin) | 0.982290673 | 0.999108571 | VR_C4 |
| A-674563 | 0.195045808 | 0.746417567 | VR_C4 |
| Dasatinib | 0.982290673 | 0.999108571 | VR_C4 |
| Doramapimod (BIRB 796) | 0.982290673 | 0.999108571 | VR_C4 |
| Dovitinib (CHIR-258) | 0.222570439 | 0.103903473 | VR_C4 |
| Foretinib (XL880) | 0.195045808 | 0.73636537 | VR_C4 |
| INK-128 | 0.982290673 | 0.999108571 | VR_C4 |
| JAK Inhibitor I | 0.982290673 | 0.999108571 | VR_C4 |
| JNJ-28312141 | 0.640935916 | 0.442281378 | VR_C4 |
| KI20227 | 0.982290673 | 0.843928758 | VR_C4 |
| PI-103 | 0.523581003 | 0.73636537 | VR_C4 |
| PP242 | 0.982290673 | 0.999108571 | VR_C4 |
| Pelitinib (EKB-569) | 0.0019955 | 0.004034813 | VR_C4 |
| Quizartinib (AC220) | 0.195045808 | 0.256332699 | VR_C4 |
| Tivozanib (AV-951) | 0.769943707 | 0.746417567 | VR_C4 |
| AT7519 | 0.982290673 | 0.999108571 | VR_C4 |
| BEZ2351 | 0.982290673 | 0.999108571 | VR_C4 |
| PD173955 | 0.391261368 | 0.552299779 | VR_C4 |
| Rapamycin | 0.982290673 | 0.999108571 | VR_C4 |

These data along with transcriptional, mutational, and phenotypic heterogeneity observed in the VR_Cs (**Figure 2**) indicate that VEN resistance is characterized by heterogeneous genetic, transcriptional, and phenotypic properties that translate to distinct therapeutic vulnerabilities.

### Validating VR_Cs by mapping VRS onto other expression cohorts

To assign VRS states to other transcriptomic datasets, we devised a PCA (Principal Component Analysis) projection approach (see **Methods** and **Figure S3A**). Briefly, genes used for NMF decomposition were used to perform PCA on all VR_C and VSC samples in the discovery cohort (BeatAML1). The expression of the same genes in the target dataset are extracted and transposed. The dot product between the transposed expression matrix and gene PC loadings from BeatAML1 gives the sample PC loadings of the target dataset, projecting them into the same PCA space as the samples from BeatAML1. K-mean clustering was then used to assign cluster definitions to the projected samples based on their distance from the centroid of the original clusters. Using this approach, we projected cluster definitions onto 3 cohorts BeatAML2, CCLE (AML cell-lines) and (The Cancer Genome Atlas) TCGA-AML (**Table 1**). In both BeatAML2 and CCLE samples classified as VR_Cs were enriched for VEN resistant samples and had higher AUC for VEN (**Figure S3B-E**). Many of the mutated genes associated with specific VR_C in BeatAML1 (**Figure 2C**), showed significant association with projected VRS definitions and conserved patterns of association with different VR_Cs (**Figure S3F-G**). Drugs identified in BeatAML1 with cluster specific sensitivity (**Figure S2M**) were recaptured at significant rates in BeatAML2 (**Figure S3H**). The projected clustering also recapitulated major cell composition (**Figure S3I-J**) and pathway enrichment (**Figure S4**) patterns in BeatAML2 and TCGA. **Supplementary Notes 1** discusses the above results in greater detail. These results indicate that the projected clustering definitions preserve characteristics of the original clustering and can facilitate further exploration.

### VRS reflect developmental expression programs coopted by AML blasts that result in VEN resistance

Some VRS showed distinctive enrichment for blasts in specific developmental compartments (**Figure 2D, S2E and S3I, J**). VR_C1 was also consistently enriched for FAB_M4/5 AMLs (**Figure 2C and S5A**). Studying the expression of hematopoietic cell-type marker genes in the original and projected VRS (**Figure S5B**), we observed: consistent over-expression of Mono, ProMono and cDC markers in VR_C1, which were in turn suppressed in VR_C4 and VSC. VR_C2 lacked a systematic pattern and showed over-expression of some markers specific for immature blasts, some cDC markers and suppression of erythroid markers. VR_C3 showed over-expression of erythroid and megakaryocytic markers. However, it’s unclear if VRS specifically capture these developmental programs. To address this, we projected VRS definitions onto bulk-RNAseq from sorted mature and stem-cell compartments of monocytic and primitive-like leukemias^19^. These samples segregated by their cellular phenotype and not the clinical classification of the patients they were derived from (**Figure S5C**). Projecting VRS definitions onto these samples, we found that VRS were not associated with the clinical phenotype of the samples (**Figure 3A**), but with their cell-type (**Figure 3B**). All mature blasts were classified as VR_C1. The leukemia stem cell-like and non-leukemia stem-cell samples were distributed across VRS. VR_C4 and VSC were predominantly (> 70%) composed of leukemia stem-cell like samples.

**Figure 3:**
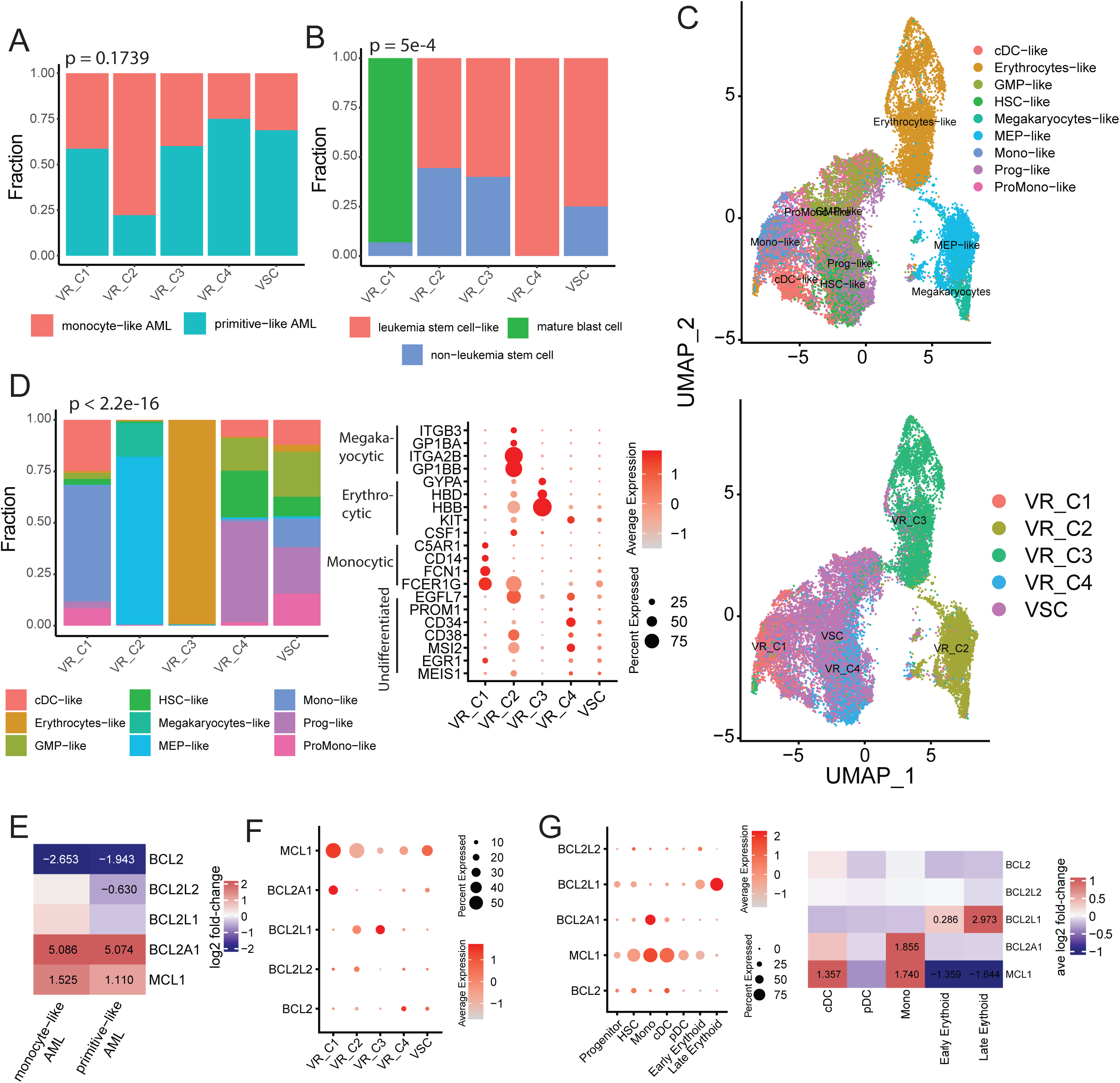
Some VR_Cs recapitulate characteristics of hematopoietic development: Fraction of clinical sample-type **(A)** and sorted cell-type **(B)** in each VRS. **C)** uMAP of AML blasts colored by their developmental status (top) and projected VRS definition (bottom). **D)** Fraction of each developmental blast category across VRS (left) and dot plot depicting expression of hematopoietic cell-type markers across VRS (right). **E)** Heatmap of log2FC of anti-apoptotic in VR_C1 relative to other samples in each clinical sub-type. Log2FC reported when significant (p.adj < 0.1). **F)** Dot plot depicting expression of anti-apoptotic genes in VRS projected onto single cells (in **C)**. **G)** Dot plot depicting expression of anti-apoptotic genes across cell-types in normal bone marrow (left) and heatmap of the average log2FC of these genes in indicated cell-type relative to primitive cell-types (HSC and Progenitor). The log2FC is reported when significant (pa.dj < 0.01 and absolute log2FC > 0.25). **Note:** association between two factor variables was tested using Chi-square test.

To explore the developmental relationships of VRS in greater granularity, we used scRNAseq data from AML blasts with developmental classifications from two studies^20,32^ covering blasts over a spectrum of developmental states (**Figure 3C**, top). Projecting VRS definitions onto these cells (**Figure 3C-D**), we found that VR_C1 was enriched for mature blasts i.e., Mono-like and cDC-like blasts (**Figure 3C-D**). VR_C2 and VR_C3 in contrast showed enrichment for megakaryocytes/MEP (megakaryocytic-erythroid progenitor)-like and erythrocyte-like blasts, respectively (**Figure 3C-D**). Intriguingly, VR_C4 were predominantly Prog-like (**Figure 3D**, left). VSC’s however showed a mixture of blast types with a combined enrichment for less developed blasts (Prog-like (22.6%), HSC-like (9.3%) and GMP-like (21.9%)). The blast-type composition of the VRS also mirrored expression of associated cell-type markers (**Figure 3D** right).

We next analyzed the expression of anti-apoptotic genes, whose over expression is a common mechanism of VEN resistance^1,5^. When we compared VR_C1 samples to all other sorted samples in monocytic-like and primitive-like AMLs independently, we observed suppression of *BCL2* and over-expression of *MCL1* and *BCL2A1* in both comparisons (**Figure 3E**). At the resolution of single cells (**Figure 3C),** VR_C1-3 were characterized by low *BCL2* expression and high expression of anti-apoptotic genes (**Figure 3F**). VR_C1 showed high expression of *MCL1* and *BCL2A1*, VR_C2 showed modest expression of *MCL1* and *BCL2L1* in a similar fraction of cells as those observed in VR_C1 and 3, respectively. VR_C3 was characterized by high expression of *BCL2L1* with low to no expression of *MCL1* and *BCL2A1*. Expression of *BCL2* was mostly localized in VR_C4 and VSC, intriguingly they also showed modest expression of *MCL1*. These patterns (**Figure 3F**) mirror expression patterns observed in normal hematopoietic cells (**Figure S5D** and **3G**) mimicked by the VRS (**Figure 3B-D**): *BCL2* (progenitor and HSC; corresponding to VR_C4 and VSC), *MCL1* (monocytes and cDC; corresponding to VR_C1), *BCL2A1* (Monocytes; corresponding to VR_C1) and *BCL2L1* (erythroid; corresponding to VR_C3).

These data indicate that some of the VR_Cs (VR_C1 and 3) capture aspects of normal hematopoietic development that in the malignant context are associated with resistance to VEN; and can do so at the resolution of bulk tumors and single cells.

### VR_C3 is sensitive to JAK inhibition and show synergy with VEN in a subset of VR_C3 like cell lines

We next focused on identifying specific vulnerabilities in VR_Cs that could be exploited to overcome VEN resistance in these states. While we found no drugs with specific sensitivity in VR_C3 across BeatAML1 and 2 (**Figure S2N** and **3H**), at the level of inhibitor families (see **Methods**, **Figure 4A**) VR_C3 showed sensitivity to several families of RTK inhibitors. Intriguingly, VR_C3 was also characterized by activation of JAK-STAT signaling across cohorts and was sensitive to JAK family of inhibitors (**Figure 2E-F**, **S4A-B** and **4A**). Across cohorts, pathways upstream (KEGG cytokine-cytokine receptor signaling) and downstream (proliferation, MAPK signaling) of JAK-STAT signaling were often significantly higher in VR_C3 relative to other VRS (**Figure 4B** and **S6A-B)**. We also observed consistent over-expression of *BCL2L1*, downstream of JAK-STAT signaling across datasets (**Figure 4C** and **S6A, C**). We further confirmed activation of JAK-STAT in VR_C3-like cell lines at the protein level using RPPA (reverse phase protein array) data (**Figure 4D**, see **Methods**).

**Figure 4:**
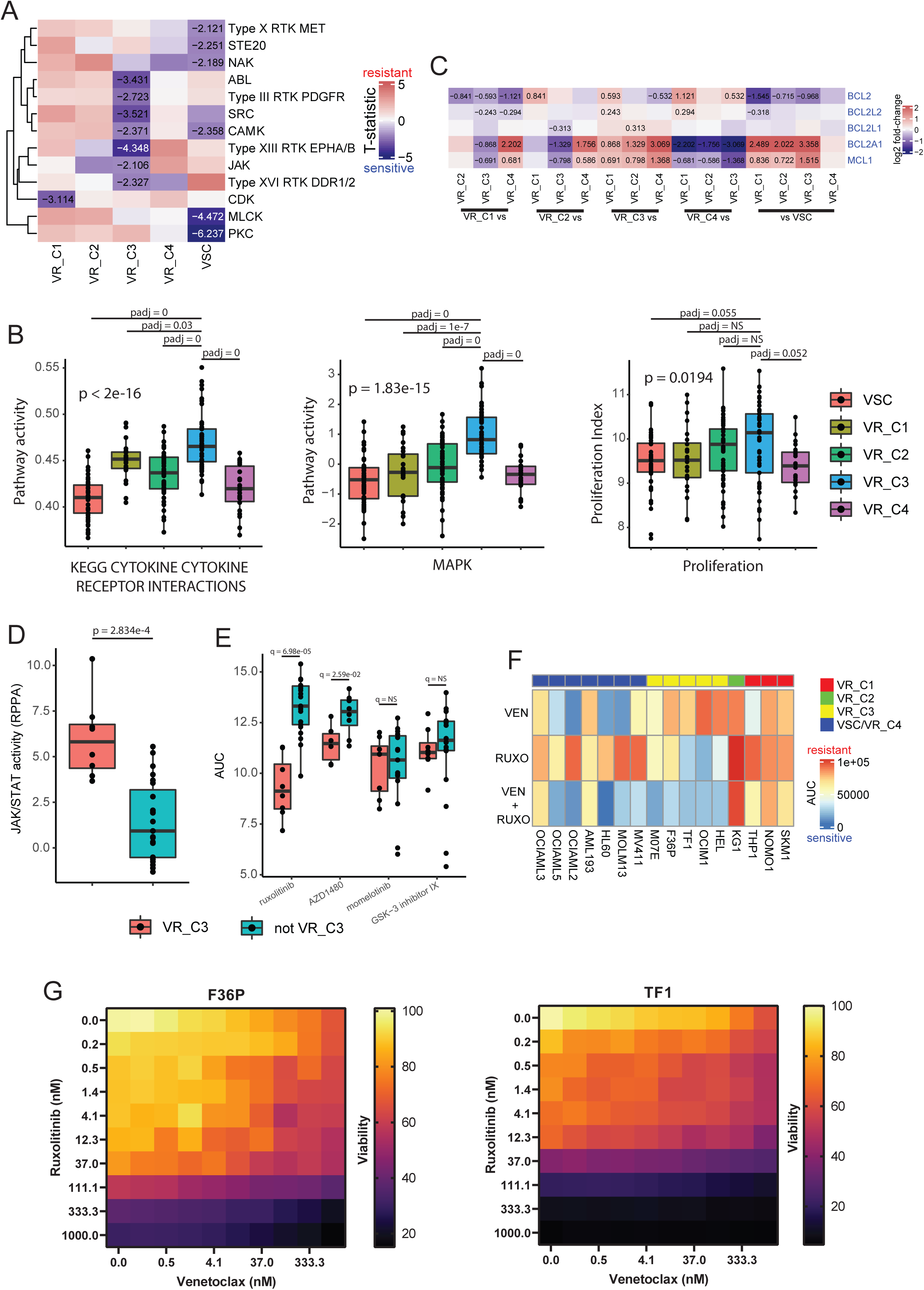
VR_C3 is sensitive to JAK-STAT inhibition: **A)** Heatmap of T-statistic comparing activity of drug family in each cluster with all other samples in BeatAML1 and 2, T-statistic is report when significantly lower (q < 0.1; one-sided T-test). **B)** Boxplot comparing pathway scores (left, mid) and proliferation index (right) in BeatAML1. P-values is computed using ANOVA, pairwise p.adj with Tukey’s test (significant at p.aj < 0.1). **C)** Heatmap of log2FC (in BeatAML1) of anti-apoptotic genes in indicated comparisons (columns) and reported when significant (p.adj < 0.1). **D)** Boxplot comparing JAK-STAT activity, inferred from protein expression (RPPA; see **Methods**; two-sided T-test). **E)** Boxplot of drug-response AUC for pan-JAK inhibitors. One-sided T-test followed by FDR correction used to test for significantly lower AUC (q<0.1) in VR_C3-like cell lines. **F)** Heatmap of drug-response AUCs for cell lines treated with VEN, RUXO and VEN+RUXO. **G)** Heatmap of viability of F36P (left) and TF1 (right) at different concentrations of combinations of VEN (x-axis) and RUXO (y-axis).

Because VR_C3 was characterized by activation of JAK-STAT signaling and pro-survival and proliferation pathways downstream of JAK-STAT (**Figure 4A-D** and **S6A-C**), we hypothesized the VR_C3 could be sensitive to inhibition of JAK-STAT signaling. In fact, VR_C3-like cell lines were significantly more sensitive to two of the four pan-JAK/JAK-STAT inhibitors tested using data from CTRP (Cancer Therapeutics Response Portal) v2^33^(**Figure 4E**). To independently test the hypothesis, we tested the response of VR_C3-like cell lines to VEN (VEN), a JAK inhibitor ruxolitinib (RUXO) and their combination: VEN+RUXO. VR_C3-like cell lines were resistant to VEN and sensitive to RUXO and VEN+RUXO (**Figure 4F** and **S6D**). Interestingly, in F36P and M07E, there was a much larger drop in AUC between treatment with RUXO and VEN+RUXO (**Figure 4F**). We therefore tested synergy between VEN and RUXO in four VR_C3 cell lines (**Figure 4G** and **S6E-F**). We detected modest synergy in F36P at most concentrations (**Figure S6F**, right) and in TF1 at low concentrations of RUXO (0.2-1 nM) and low to moderate concentrations of VEN (0.2-37 nM) (**Figure S6F**, left). Sensitivity in HEL and OCIM1 was primarily driven by RUXO (**Figure S5E**).

These results indicate that VR_C3 like cells/samples have activation of JAK-STAT signaling, consequently are sensitive to its inhibition. In a subset of cases, JAK inhibition also broke VEN resistance resulting in synergistic effects.

### VR_C1 is characterized by transcriptional signature of metabolic fitness and sensitivity to CDK inhibition

VR_C1 showed consistent over-expression of various metabolic pathways (**Figure 2E-F** and **S4A-B**) and over-expression of genes/enzymes regulating glycolysis, TCA (tricarboxylic acid) cycle and ETC (electron transport chain) (**Figure S7A**). We also observed consistent over-expression of nutrient transporter pathways in VR_C1 (**Figure 5A** and **S7B**). Taken together, this indicated that VR_C1 shows higher transcriptional activity of metabolic and nutrient uptake pathways, suggesting they might have higher metabolic fitness. VR_C1 also showed activation of PI3K-mTOR signaling (**Figure 2E-F**, S**2H** and **S4A-B**). The activity of PI3K-mTOR signaling, a known regulator of metabolic activity^28^ was positive correlation with activity of metabolic and nutrient uptake pathways (**Figure 5B** and **S7C**). Further, the activation of PI3K-mTOR signaling and metabolic pathways (glycolysis, OxPhos and FAM) were associated with VEN resistance and sensitivity to drugs that showed specific sensitivity in VR_C1 (**Figure 5C**, **S7D and S2N**). We therefore hypothesized that high PI3K-mTOR signaling in VR_C1 increases nutrient uptake and metabolic fitness resulting in VEN resistance. VR_C1 patients might therefore be sensitive to inhibition of PI3K-mTOR signaling. In fact, in BeatAML1 and 2, VR_C1 showed increased sensitivity to several PI3K-mTOR pathway inhibitors (**Figure S2N** and **Table 2**) including rapamycin (**Figure S8A**, left). In BeatAML1, VR_C1 had lowest AUC for rapamycin that is significantly lower relative to VSC and is significantly lower relative to all other VRS in BeatAML2 (**Figure S8A**, left). We also observed in both BeatAML1 and 2 a specific sensitivity to panobinostat (PANO) a HDAC inhibitor (**Figure S2N, S8A** middle; **Table 2**).To access the possible mechanism of sensitivity to PANO, we compared published bulk-RNAseq (see **Methods**) of leukemia cells harvested from a preclinical mouse model of t(8;21) AML treated with PANO^34^. PANO treatment resulted in suppressed cell-cycle pathways (G2M checkpoint, Mitotic spindle), MTORC1 signaling, MYC targets and OxPhos (**Figure S8B**), indicating that PANO might inhibit metabolism and mTOR signaling contributing to VR_C1 sensitivity to PANO. We, however, were unable to recapture VR_C1’s sensitivities to PANO, PANO+VEN (**Figure S8C**) and mTOR inhibition (**Figure S8D**) in cell line models. This inconsistency could possibly arise due to differences in metabolic properties of cell-lines and primary AMLs.

**Figure 5:**
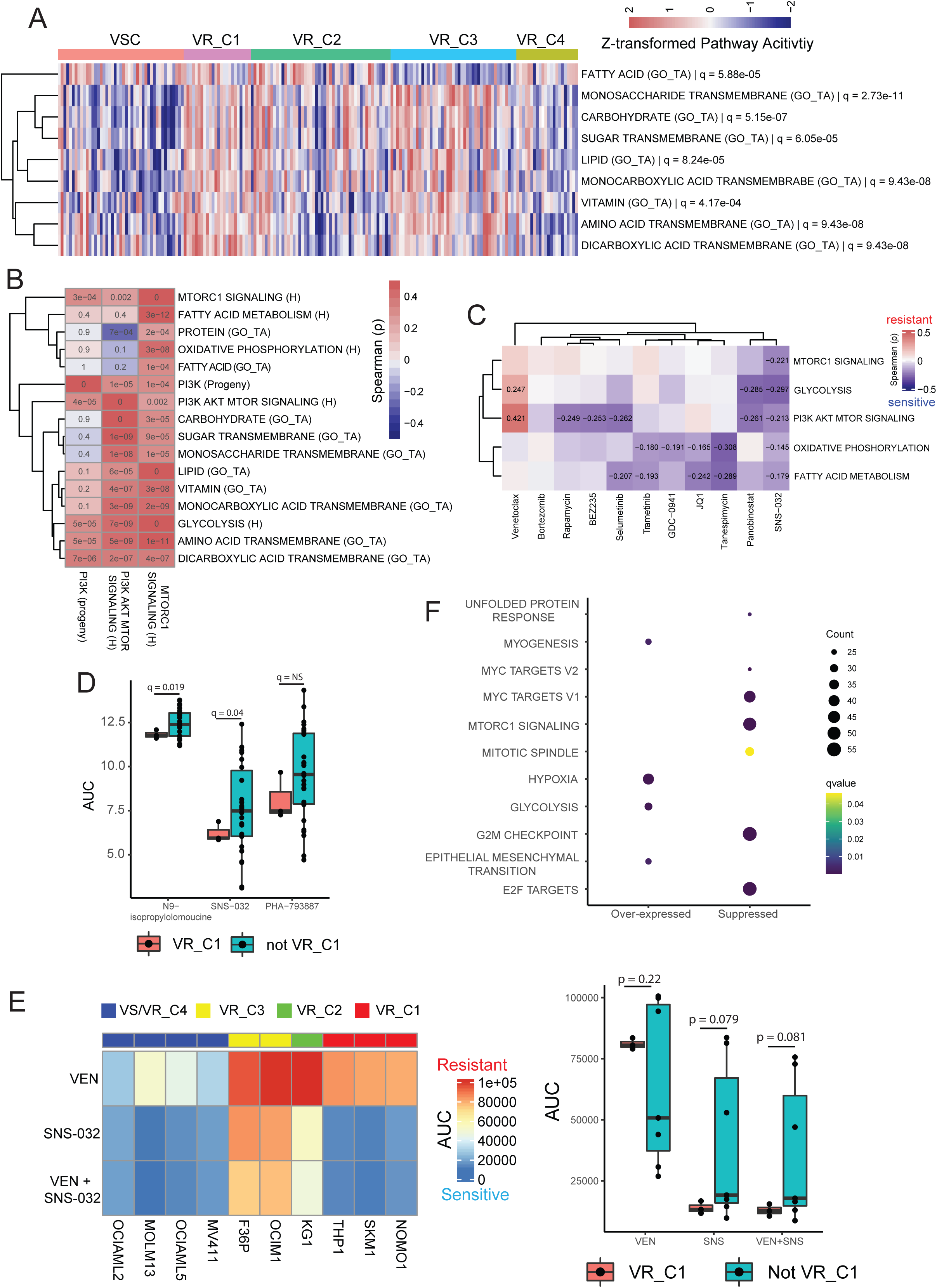
VR_C1 is characterized by transcriptional signature of higher metabolic activity and sensitivity to CDK inhibition: **A)** Pathway activity of nutrient transport pathways in BeatAML1 with differential activity across VRS (ANOVA followed by FDR correction, significant at q < 0.1). **B)** Heatmap of spearman correlation coefficients between metabolic pathways (rows) and the PI3K-mTOR signaling axis (columns) in BeatAML1. q-values after FDR correction across each column reported in cells, (significance at q < 0.1 **C)** Heatmap of spearman correlation coefficients between activity of metabolic pathways (rows) and drug-response AUCs in BeatAML1. p-values corrected by FDR for each drug (significant q < 0.1) **D)** Boxplot of drug-response AUC for CDK inhibitors. One-sided T-test followed by FDR correction used to test for significantly lower AUC (q<0.1) in VR_C1-like cell lines **E)** Heatmap of drug-response AUCs for cell lines treated with VEN, SNS-032 and VEN+SNS-032 (left) and corresponding boxplots (right) (two-sided T-test). **F)** Functions enriched (p < 0.01 and q < 0.05) of genes over and under-expressed (see **Methods**) in SNS-032 treated MOLT4 cells relative to vehicle (DMSO) treated cells. **Note:** GO_TA (gene ontology transport activity), H (Hallmark), Progeny indicate the pathway dataset used to compute pathway activity (Figure 5A).

VR_C1 was also sensitive to the pan-CDK inhibitor SNS-032 (**Figure S2N, S8A** right; **Table 2**) and showed specific sensitivity to CDK inhibitor family (**Figure 4A**). VR_C1-like cell-lines were sensitive to two of three pan-CDK inhibitors tested from CTRPv2^33^ **Figure 5D**). Testing sensitivity to SNS-032 and its combination with VEN, we recaptured sensitivity of VR_C1 cell-lines to both, though at a nominal degree of significance (p = 0.079 and p = 0.081 respectively; **Figure 5E**), likely due to the small number of VR_C1 cell-lines (n = 3). VR_C1-like cell-lines had similar sensitivity to SNS-032 and VEN+SNS-032 suggesting lack of any additive or synergistic effect between the drugs (**Figure 5E**). To gain mechanistic understanding of VR_C1’s sensitivity to SNS-032, we studied the pathways perturbed by SNS-032 treatment using bulk-RNAseq from MOLT4 cell-line treated with SNS-032 and DMSO^35^. SNS-032 suppressed genes (q < 0.1 and log2FC < -1.5) in E2F and MYC targets, MTORC1 signaling and cell cycle pathways (mitotic spindle and G2M checkpoint; **Figure 5F**). GSEA analysis, although at a lower statistical threshold (q = 0.2), also showed suppression of OxPhos along with other pathways (**Figure S8E**), indicating the sensitivity of VR_C1 to SNS-032 in part could be explained by inhibition of metabolic pathways and suppression of mTOR signaling (mTORC1 signaling; **Figure 5F** and **S8E**).

The above data therefore indicates that VR_C1 is characterized by a transcriptional activation of metabolic pathways and PI3K-mTOR signaling constitutes a key regulator of these pathways. VR_C1 also shows sensitivity to CDK inhibition, which may mechanistically function, in part, by suppressing mTOR signaling and OxPhos.

### VRS capture intra tumor heterogeneity and dynamics of cell-states in development of VEN resistance

To assess whether VRS could elucidate intra-tumor heterogeneity of cell-states associated with VEN resistance, we analyzed scRNAseq from pre- and post-treatment cells from a patient who relapsed under VEN+AZA treatment, showing expansion of a monocytic clone with NRAS Q61K and loss of a primitive SMC1A R807H clone upon relapse^14^ (**Figure 6A**, left). The cell-types of the cells were annotated by transferring labels from healthy BM (see **Methods**; **Figure 6A**, middle) and VRS definitions were projected onto cells in the myeloid compartment (**Figure 6A**, right). Interestingly multiple VRS subtypes (VR_C1, C2 and VSC) were well represented in the tumor suggesting that the tumor was a mixture of subtypes. The subtype representation in bulk data likely represents a dominance of one cell subtype. The cell subtype composition matched expectations (**Figure 2D**), with VSC cells enriched for HSCs, VR_C1 for Mono cells and VR_C2 showed a mixture of Mono and GMP cells (**Figure 6B**). Pre-treatment cells existed in both sensitive (VSC, 41%) and resistant (VR_C1 and C2, 59%) VRS (**Figure 6C**), however post-treatment VSC cells were eliminated, and resistant states came to dominate (VR_C1: 5% and VR_C2: 93%). VR_C2 showed a higher proliferation-index (see **Methods**) before and after treatment, which might provide a competitive advantage allowing it to outgrow VR_C1, despite the shared resistance to VEN (**Figure 6D**). At the pathway-level VR_C1 was characterized by higher expression of several inflammatory and metabolic pathways as well as high PI3K-mTOR and MTORC1 signaling (**Figure 6E**) consistent with earlier observations (**Figure 2E-F**). VR_C2 showed increased expression of G2M checkpoint and E2F targets (**Figure 6E**) indicating higher cell-cycle activity which is consistent with the higher inferred proliferation rates observed (**Figure 6D**). Interestingly, the VSC cells showed high expression of MYC targets and transcriptional indicators of DNA damage (DNA repair and UV response down; **Figure 6E**). These data illustrated that the VRS can be used to gain insights into intra-tumor heterogeneity and dynamics of VEN resistance associated cells states.

**Figure 6:**
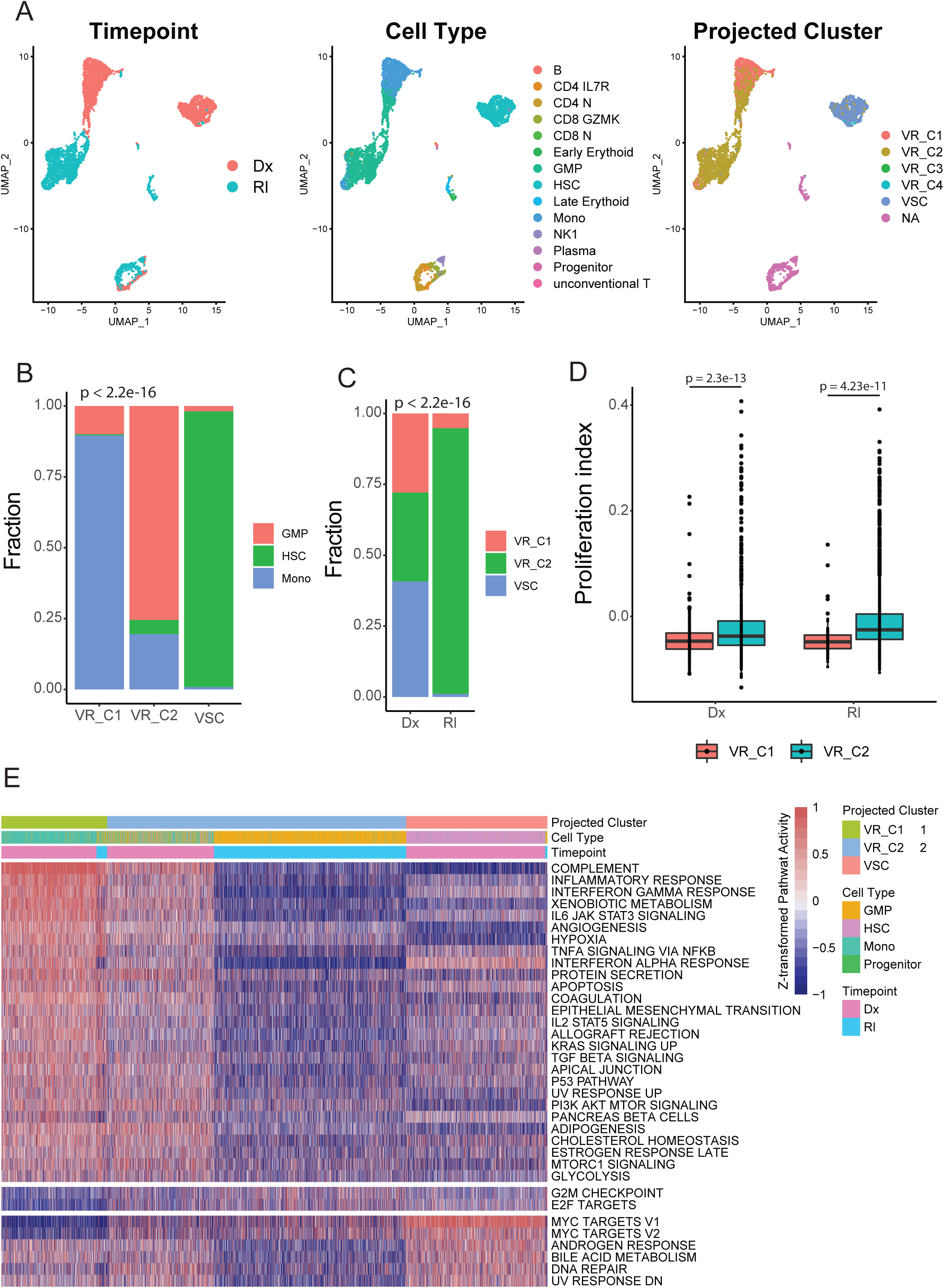
VRS facilitates interpretation of cellular dynamics in response to VEN treatment: **A)** uMAPs of cells colored by treatment time point (left), cell-type (mid) and VRS definition projected onto myeloid cells (right). **B)** Fraction of cell-types in each VRS. **C)** Fraction of VRS in pre and post treatment cells. **D)** Boxplot comparing proliferation index between VR_C1 and VR_C2 like cells pre- and post-treatment (p-values computed using two-sided T-test). **E)** Heatmap of pathway activity in cells for Pathways over-expressed (log2FC > 0.15 and p.adj < 0.01) in VR_C1, VR_C2 and VSC. **Note:** association between two factor variables was tested using Chi-square test.

## Discussion

VEN in combination with hypomethylation agents has emerged as a paradigm shifting treatment for AML, however primary and acquired resistance^4,5^ limit the long-term efficacy of therapy. While several mechanisms for VEN resistance have been identified^1,5,8–14,16,21^, the heterogeneity of these mechanisms in VEN resistant patients has not been fully elucidated. In this study, we decomposed expression of genes associated with VEN resistance to identify four transcriptionally distinct VEN resistant patient groups. These groups are characterized by distinct phenotypic, mutational and drug response characteristics. Elucidating such heterogeneity can facilitate identification of specific vulnerabilities in VEN resistant patients and improve selection of appropriate therapies specific to each patient.

VEN resistance has been linked to differentiation status of AML blasts, with monocytic tumors being resistant to VEN-based therapies^14–16,21,36^. We recapture this association in VR_C1 patients. Further using bulk-RNAseq from sorted cell-types and scRNAseq, we show that the VR_C1-like expression state is associated with monocytic differentiation at a more granular resolution. In contrast, VR_C3 showed a strong preference for erythroid-like blasts. VR_C1 and VR_C3 showed suppression of expression of *BCL2* and over-expression of other anti-apoptotic genes such as *MCL1* (VR_C1), *BCL2A1* (VR_C1) and *BCL2L1* (VR_C3), rendering them inherently resistant to VEN. Association of erythrocytic differential and VEN resistance has been suggested by Kuusanmaki et al. ^20^, stating that FAB-M6 and M7 AMLs are resistant to VEN in a BCL2L1-dependent manner. This mechanism of resistance may however be more widespread, as we identify VR_C3-like tumors outside FAB-M6/7.

Inhibiting anti-apoptotic genes over-expressed in VR_C1 and VR_C3 presents a potential therapeutic strategy to treat VEN resistant monocytic and erythroid like tumors. While these strategies have been effective *in vitro*, toxicity associated with these inhibitors^22,23^ has limited their clinical application. Further, VR_C3-like tumors showed an enrichment for TP53 mutations which are commonly associated with poor outcome and resistance to chemotherapy^37,38^. Therefore, identifying alternative strategies to treat these VEN resistance AML subtypes can have a significant clinical impact.

Transcriptional characterization of VR_C1 indicates a transcriptional activation of metabolic pathways, likely mediated by active PI3K-mTOR signaling, resulting in improved metabolic fitness. Improved metabolic fitness has been linked to VEN resistance^10^. mTOR signaling is a core regulator of metabolism^28^and its over-expression can drive the observed activation of metabolic pathways. Consistent with these findings, VR_C1 is characterized by sensitive to mTOR inhibition. Additionally, VR_C1 is sensitive to CDK inhibitors, like SNS-032, which suppressed OxPhos and mTOR signaling. Intriguingly, mTOR inhibition was ineffective in cell line models, likely because they don’t recapture metabolic properties of patient derived cells. SNS-032 was however effective in both models, indicating its effects are multifactorial including, but not limited to mTOR inhibition. Taken together, these findings warrant further exploration of PI3K-mTOR and CDK inhibition as therapeutic strategies to treat Mono-like AMLs.

VR_C3 was characterized by activation of JAK-STAT signaling, down-stream activation of pro-survival pathways, proliferation, and consequent sensitive to JAK inhibition. Intriguingly in a subset of VR_C3-like cell-lines, JAK inhibition by RUXO broke VEN resistance, resulting in synergy for the combination. This observation suggests that treatment with RUXO alone or in combination with VEN might be effective in VR_C3-like AMLs, which warrants further exploration. VR_C3 was also characterized by higher infiltration by CTLs, high expression of cytotoxic immune effector and immune checkpoints genes, suggesting that VR_C3 AML may also benifit for immunotherapy.

Consistent pathway activation relative to other VRS was absent in VR_C2. We however observed consistent suppression of *HOX* expression and transcriptional activity. This was consistent with absence of NPM1 mutations in VR_C2^39^. Suppression of HOX has been previously reported in VEN resistant samples^30^. However, it is unclear if it contributes mechanistically to VEN resistance or is simply a consequence of the absence of NPM1 mutations. VR_C2 is also characterized by enrichment of RAS mutation (especially NRAS), which have been associated with VEN resistance^14,21^. As expansion of RAS clones have been linked to relapse under VEN-based therapy^14^, establishing a link between prevalence of RAS mutation in a tumor and loss of *HOX* expression can facilitate development of diagnostic approaches that can detect relapse under VEN-based therapy associated with RAS mutations.

Though VR_C4 consisted of VEN resistant patients, they were enriched for less differentiated blast and shared transcriptional characteristics with VSC. Mutationally VR_C4 like VSC had high mutation rates in *FLT3* and *NPM1*, but also in *DNMT3A* unlike VSC. Mutations in *DNMT3A* may be associated with better response to AZA and cytarabine^40,41^. Thus VR_C4-like tumors may still benefit from current VEN combination therapies and perhaps with some selectively targeting DNMT3 inhibitors^42^.

Projecting VRS definitions onto single cell profiles from a patient who relapsed under VEN+AZA, we identified co-existence of multiple VR_Cs and how their proportions changed after therapy. Indicating VRS can also capture intra-tumor heterogeneity in the context of VEN resistance. Elucidating VEN resistance associated intra-tumor heterogeneity can facilitate development of effective combination therapies.

Taken together, in this study we capture transcriptomic heterogeneity in VEN resistant patients, which translates to distinct phenotypic, mutational and drug response profiles (**Figure 1-3**). This additional granularity facilitated identification of groups of patients with different mechanisms of VEN resistance and their specific vulnerabilities, that can be targeted with VEN or selective combinations (**Figure 4-5**). We also show that multiple VRS can co-exist in a single tumor, therefore also capturing intra-tumor heterogeneity (**Figure 6**)

## Limitations of study

Our analysis of heterogeneity in VEN resistant patients utilizes bulk-RNAseq and *ex-vivo* drug response from BeatAML^15,26^. VEN is commonly used in the clinical setting in combination with AZA or cytarabine. It is therefore likely that some of these observations might not directly translate to clinical cohorts or settings. Further, responses *in vitro* may not always translate to responses in patients, underlining the need for further analysis with data from patients treated with VEN combinations. Recent studies have also suggested that studying sub-population of LSCs can be critical to understanding the development of resistance and relapse to VEN therapies in AML^19,43^. It is challenging to capture this level of granularity using bulk expression data and will require single cell profiling of the LSC compartment. In **Figure 3**, we capture developmental preferences in VRS at the resolution of single cells. VR_C3 and VR_C2 showed preference for erythrocyte-like and MEP-like blasts. These cells however came from two previously reported^20^ samples, respectively. While we do observe over-expression of erythroid markers in VR_C3 across bulk cohorts (**Figure S5B**), the single cell patterns should be interpreted with caution and warrant further analysis in large cohorts.

VR_C1 shows elevated expression of metabolic pathways. While pathway analysis can capture basic trends of metabolic activity, precise changes in metabolic pathways and utilization of metabolites can be challenging to infer from gene expression. It is likely that through detailed metabolomic characterization VR_C1 can be further sub-divided based on metabolic pathways used to meet their energetic requirements. Another limitation in this scenario is using cell lines for downstream validation. Metabolism of cell lines can be sensitive to culture conditions that can differ across cell lines. As our analysis requires comparisons across diverse set of cell lines instead of comparisons between parental and modified cell lines, it is challenging to validate metabolic patterns in VR_C1 using cell line models.

## Star Methods

### Datasets

mRNA expression (counts), mutation calls and drug response (AUCs) data for the first release of BeatAML (BeatAML1) was obtained from Tyner et al.^26^ using the data portal http://www.vizome.org. mRNA expression (counts), mutation call, drug response (AUCs) for the second release of BeatAML^15^ (BeatAML2) as well as inhibitor family information and clinical data for all patients was obtained from https://biodev.github.io/BeatAML2/. mRNA expression (log2TPM+1) and protein expression (RPPA) for CCLE^44^ cell lines was downloaded from (20Q4: https://depmap.org/portal/) and (https://data.broadinstitute.org/ccle/) respective. Drug response data (AUCs) for CCLE cell lines was obtained from CTRPv2^33^. mRNA count data for AML TCGA^45^ samples was obtained from firehose (http://firebrowse.org/), normalized mRNA counts, mutation and clinical data was obtained from UCSC Xena^46^ (https://xenabrowser.net/datapages/). mRNA counts of leukemia cells from mice treated with PANO and vehicle control was obtained from GSE198119^34^. Whole transcriptome differential expression statistics (log fold changes and q-values) comparing MOLT4 cell line treated with SNS-032 to DMSO treatment was obtained from GSE206612^35^. FASTQ file of bulk-RNAseq from sorted sub-populations of AML blasts were obtained from Waclawiczek et al.^19^.

scRNAseq counts (10x genomics) for normal bone marrow were obtained from Abbas et al.^47^ CITE-seq for VEN+AZA treated patient was obtained from GSE143363^14^. scRNAseq counts for AML blasts was obtained from Van Galen et al.^32^ (SeqWell) and Kuusanmaki et al.^20^ (10x genomics).

Mouse and Human Hallmark^48^ gene sets were obtained from MsigDB^27^. Gene-sets for nutrient transporters were obtained from GO molecular function^49^ in MSigDB. Through manual inspection we select pathways that define transporters for amino acids, carbohydrates, fatty acids, lipids, proteins, sugars, monosaccharides, and vitamins.

### Cell culture

The AML cell lines: MV4-11, MOLM13, HL60, AML193, SKM1, NOMO1, THP1, OCIAML2, OCIAML3, OCIAML5, HEL, OCIM1, TF-1, F36P, MO7E and KG,1 were obtained from ATCC and DSMZ, and authenticated by short tandem repeat DNA fingerprinting. Cell lines were maintained if not otherwise specified in RPMI-1640 medium (SIGMA) with 2 mM Glutamine (SIGMA) containing 10–20% heat inactivated (h.i.) fetal calf serum (FCS) and 1% penicillin-streptomycin (Life Technologies Laboratories), as suggested by cell lines provider. Additionally, F36P, MO7E, OCIM1, OCIAML5, and TF1 were supplemented with 10 ng/ml GM-CSF, and AML193 was cultured in IMDM with 30% h.i. FCS and supplemented with IL3 and GM-CSF at 10 ng/ml each. All cells were grown at 37 °C in a humidified atmosphere containing 5% carbon dioxide and passaged every 2-3 days at the density of 0.5 million of cells/ml.

### Identifying resistant and sensitive cell-lines/patients

For a drug, high AUC values indicate resistance and low values sensitivity. AUC values of each drug in either CCLE (AML and CLL) cell lines or in BeatAML samples were modeled using Gaussian mixture models, fitting up to 3 normal distributions to the data using the *flexmix* R package (https://cran.r-project.org/web/packages/flexmix/index.html). The optimal model was selected using the “ICL” option in the function *getModel()*. If 2 or 3 normal distributions fit the AUC data and the difference between means of the distributions with highest (HDis) and lowest (LDis) means was greater than the sum of their standard deviations and mean (LDis) < 10^th^ %ile of AUCs in the dataset. Samples that were part of LDis were considered sensitive to the drug and those in HDis were considered resistant. In all other cases a hard threshold of AUCs < 10^th^ %ile across all AUCs in the dataset was used to delineate sensitive samples from resistant. A threshold of 10 AUC was used for CCLE (**Figure S1A**), and 112 AUC was used for BeatAML.

### Drug response AUC and synergy calculation

Selected cell lines were plated at a density of 10,000-20,000 cells/well in complete RPMI-1640 (SIGMA) supplemented with 10-20% fetal calf serum onto 96 well plates and subjected to treatment with Ruxolitinib (Selleckchem), Venetoclax (ABT-199) (Selleckchem), Panobinostat (Selleckchem) and CDK2,7,9 inhibitor SNS-032 either as monotherapy or as combination of Ruxolitinib/VEN or Panobinostat/VEN, or SNS-032/VEN for 72 h. In one experiment, each treatment condition was seeded in triplicates, each experiment was performed independently 3–5 times.

Following drug concentrations were used: VEN was used at doses ranging from 0 to 1000 nM for 72 h, Ruxolitinib at doses ranging from 0 to 1000 nM for 72 h, SNS-032 at doses ranging from 0 to 1000 nM for 72 h, and Panobinostat at doses ranging from 0 to 100 nM.

Viable cell numbers were measured by quantifying ATP using a CellTiter-Glo Luminescent Cell Viability Assay (Promega).

The response from each technical replicate was calculated. Mean values summarized from responses from each independent experiment. Dose-response curves were analyzed using a curve fitting routine based on nonlinear regression. The same range of 10 drug concentrations was tested for each cell line to compute a curve-free area under the dose–response curve (AUC) based on linear interpolation of the 10 data points using a baseline of 0 (=100% inhibition) and a maximum of 100 using GraphPad Prism ver. 9^50^. The AUCs generated for each cell line were further summarized based on VEN resistance status and Clustering Status.

Additionally, to evaluate the synergistic effects of Ruxolitinib and VEN, cells were seeded in 96-well plates at densities as described above, creating 10 × 10 matrix of treatment. Viable cell numbers were measured by quantifying ATP using a CellTiter-Glo Luminescent Cell Viability Assay (Promega). The results of five independent experiments were collected, processed, and visualized for synergy evaluation using COMBENEFIT software^51^.

### Differential gene expression and GSEA pathway analysis

Differential expression analysis was performed using count data as an input to *DEseq2*^52^. GSEA was performed using gene lists sorted by log2 fold change (log2FC) using the *gag*e^53^ and Hallmark pathway definitions. Differentially expressed pathways are identified at q-value < 0.1. Stat.mean output by *gage* captures magnitude and direction of dysregulation of pathways, positive values indicate over-expression and negative suppression.

### Gene set over-representation analysis

Given a list of genes, analysis of over-represented pathways in them was performed using the *compareCluster()* function implemented in the Bioconductor package *clusterProfiler*^54^ with option *fun = enricher*. Over-represented pathways are identified at p-value < 0.05 and q-value < 0.1, unless specified otherwise.

In case of SNS-032 treated cells compared to vehicle treatment DEGs were defined at q < 0.1 and log2FC > 1.5 and < -1.5 for over-expressed and suppressed genes, respectively. These genes were used as the input for over-representation analysis. Significant Hallmark pathways were identified at p-value < 0.01 and q-value < 0.05.

### Single sample pathway and signaling activity quantification from bulk gene expression

Normalized gene expression was converted to Hallmark pathway activity using single sample GSEA (ssGSEA) implemented in *GSVA*^55^.

Activity of 14 signaling pathways were quantified using *Progeny*^29^ using top 100 genes associated with the activation of respective pathways.

### Deriving a transcriptional signature associated with VEN resistance

A transcriptional signature for VEN resistance was derived from AML and CML CCLE cell lines. First resistant and sensitive cells were identified, and DEGs (absolute log2FC > 1 and adjusted p-value < 0.1) between them were identified using DEseq2^52^ while controlling for tumor type (**Figure 1A**).

The log2 normalized counts of these genes in the BeatAML discovery^26^ cohort (BeatAML1) were extracted and genes which mean expression < 1 and variance < 2 were filtered out. The resultant matrix was decomposed using NMF (see below).

### Identifying clusters of VEN resistant patient by decomposing the VEN expression signature

Gene expression of the VEN resistance signature in VEN resistant samples(**M**) in the BeatAML1cohort was decomposed (**Figure 1B**) with NMF (non-negative matrix factorization) using the R package *NMF*^56^ with the options *seed* = "ica" and *nrun* = 100. NMF decomposition was performed with the number of NMF components (**K**) ranging from 2 to 10. To select the optimal number of components we adopted the approach described in Frigyesi et al.^57^. A random expression matrix (**M_R_**) was generated from the original matrix using the *randomize()* function in the *NMF* package, and the NMF decomposition procedure was repeated. The final rank for NMF is selected as the largest value of **K** for which decrease in RSS (residual sum of squares) from K-1 to K is greater in **M** relative to **M_R._** (**Figure S1D**, **K**= 6 was selected for the final NMF, resulting in 6 NMF components: C1-C6).

The patient loading of the NMF components was rescaled to range from 0 to 1. Functional correlates of these components were identified by correlating the loadings with Hallmark ssGSEA pathway activity using spearman correlation. C1, 2 and 5 showed strong correlation between their loadings and were also associated with similar pathways (**Figure S1E-F**). C2 and 5 were thus discarded. The components C1, 3, 4 and 6 were then clustered using consensus clustering (**Figure 1C**) implemented in the R package *ConsensusClusterPlus*^58^. Optimal number of clusters were identified using the procedure described in Senbabaoglu et al.^59^. The procedure yielded 4 clusters (VR_C1-4) of VEN resistant patients (**Figure 1**).

### Associating clinical variables with patient clusters

Clinical variables were divided into 1. Factor variables and 2. Continuous variables. Association of factor variables with VRS was tested using Chi-square test and ANOVA was used in the case of continuous variables. FDR correction was performed in both cases separately and significant variables were identified at q < 0.1. In the case of continuous variables that reached significance pairwise differences between groups were evaluated using Tukey’s test.

### Mutational analysis

Across datasets (TCGA, BeatAML1 and 2), the mutational status of genes was binarized, 1-mutant and 0 – wild-type. Association between mutation status of a gene and VRS was tested with Chi-square test. P-values were corrected for multiple testing with FDR and significant mutations were identified at q < 0.1. The testing was limited to genes mutated in greater than 5% of patients in the BeatAML1 (**Figure 2C**). Testing in the validation cohorts (BeatAML2 and TCGA) was limited to significant genes identified in BeatAML1.

Mutational mutual exclusivity analysis was performed at the level of each VRS in BeatAML1 using the function *pairwise.discover.test*() implemented in the package DISCOVER^60^ and was limited to genes used for association testing (see above). Mutually exclusive gene pairs were identified at q-value < 0.1.

### Proliferation Index

Proliferation rate was inferred from RNAseq data using the R library *ProliferativeIndex* (https://cran.r-project.org/web/packages/ProliferativeIndex/vignettes/ProliferativeIndexVignette.html). Variance stabilized normalized read counts obtained from the *varianceStabilizingTransformation()* function in DEseq2 was used as input to the function *ProliferationIndex()* to compute proliferation rate in each sample. Proliferation score in case of single cell data was computed using the MetaPCNA signature^61^ and *AddModuleScore()* function in Seurat^62^.

### Quantifying activity of transcription factors from gene expression

Activity of transcription factors (TFs) in each sample was estimated from normalized expression in the BeatAML1, BeatAML2 and TCGA using VIPER^63^ while controlling for pleiotropic regulation. An AML specific regulatory network was constructed from TCGA expression data using ARACNE^64^. Differential activity of TFs between two groups is tested using the function *rowTtest*(), p-values are corrected with FDR. For each TF i the difference in mean activity between groups is Mdiff_i_. A TF i is called differentially expressed if

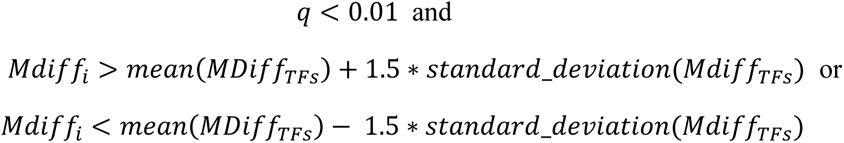

Where Mdiff_TFs_ the vector of difference in mean activity of all TFs tested. This approach was used to identify differentially active TFs in VR_C2 relative to other VRS.

### CIBERSORTx cell fraction estimation from gene expression

Cell type abundance was estimated from bulk gene expression profiles of samples BeatAML2 (including samples from BeatAML1) and TCGA using CIBERSORTx^65^ web-portal (https://cibersortx.stanford.edu/index.php). Single cell expression counts from Van Galen et al.^32^ were used as reference samples to create a signature matrix. Deconvolution for each dataset was performed independently. For each dataset gene expression data in CPM (counts per million reads) was used as input. CIBERSORTx “Impute Cell Fractions” was run in absolute mode, using S-mode for batch correction, no quantile normalization and 100 permutations. To quantify relative abundance of leukemia cells, the abundance scores of six leukemia populations in each sample were normalized to sum to 1.

### Identifying drugs, a group of samples are sensitive to

For each VR_C in BeatAML1 drugs they are sensitive to were identified by 1. One-sided T-test to test whether AUC value of the drugs was lesser in the cluster relative to all other samples and 2. Whether the cluster was enriched for samples that are sensitive to the drug, tested using the hypergeometric text. P-values in both cases are corrected for multiple testing using FDR. Significant drugs are identified at q < 0.1 for both approaches and merged. A similar approach is used for identifying drugs that all VRP are sensitive to as a collective. The testing is limited to drugs that at least 10% of the resistant samples are sensitive to.

In case of projected VRS definitions (see below) in BeaAML2 the same approach as above is used to identify drugs that each VR_C is sensitive to. The testing is however limited to drugs identified in BeatAML1 for each cluster and the fraction of drugs recaptured is calculated. To quantify the significance of the overlap we shuffle VRS membership of samples in BeatAML2, and cluster specific drugs are re-identified and overlap rate for each VR_C with corresponding VR_C specific drugs identified in BeatAML1 is computed. The procedure is repeated 5000 times. A p-value is computed for each VR_C as the fraction of times a randomly generated VRS definition has a greater overlap than the true projected VRS definitions.

### Projecting cluster definitions onto new expression data

To project VRS definitions onto new samples, we extracted the expression matrix of the genes used to perform the original NMF with gene expression scaled and centered from BeatAML1 (resistant and sensitive samples; Re) and the target dataset (Te). Genes common to both datasets are retained. PCA (Principal component analysis) analysis is performed using Re generating sample PC loadings (rPC) and gene PC (Principal components) loadings (gPC). i^th^ co-ordinate of the centroid in PCA space for each VRS are computed:

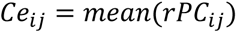

Where Ce_ij_ is the i^th^ coordinate for centroid of the VRS j and rPCij is the vector of PCA loadings of the ith PC in samples of VRS j. The PCA loadings of the target dataset (tPC) is computed by taking the dot product between the transpose of Te with gPC. The analysis uses 40 PCs, which accounts for 75% of the variability in Re.

With samples from the discovery and target dataset in the same PCA space we use K-means clustering to assign each sample in the target dataset to a VRS by passing the PCA loadings of the target samples (tPC) and coordinates of the centroids of VRS from the discovery cohort (BeatAML1) to the *kmeans* function in R. The approach is graphically summarized in **Figure S3A**. This approach was used to project VRS definitions onto samples from BeatAML2, TCGA, CCLE and sorted AML sub-compartments^19^. In case of BeatAML2 samples that overlapped with BeatAML1 were excluded. We also projected VRS definitions onto malignant cells (single cells) from VanGalen et al.^32^ and Kuusanmaki et al.^20^, and the myeloid compartment of single cell profiled in a patient who relapsed under VEN+AZA therapy from Pie et al.^14^ using their normalized expression as input.

### Response to drug families

The response profile to a family of drugs was quantified using the approach described in Bottomly et al. ^15^, briefly drug response AUC values are rescaled to range between 0 and 1. The drug scores for drugs in each family are merged using ssGSEA implemented in GSVA^55^ without normalizing the results. The transformation was performed using all the samples in BeatAML2 (including samples from BeatAML1). To identify drugs families each VRS is sensitive to one-side T-test was used to test whether the drug family scores in the VRS of interest were lower relative to all other samples. P-values were corrected for multiple testing using FDR and drugs were identified at q < 0.1. The testing was performed independently for each VRS. VRS definitions of samples from BeatAML1 (discovery) and BeatAML2 (projected) were combined for the analysis.

### CCLE RPPA JAK/STAT activity

Activity of JAK/STAT pathway was defined as the sum of expression of JAK2, STAT3, STAT3_pY705 and STAT5. Differential activity between groups was tested using a two-sided T-test.

### Analysis of gene expression in flow sorted AML blasts

Gene expression was quantified from fastq files using RSEM^66^, quantification was performed using *rsem-calculate-expression* with the Bowtie2^67^ aligner and the hg19 reference genome. Gene expression counts and TPM were read in from the RSEM outputs using tximport^68^. TPM values were transformed as log2(TPM+1) and PCA was performed using the top 5000 variably expressed genes.

Differential expression was performed comparing VR_C1 like samples to other samples in monocytic-like and primitive-like AML patients separately using DESeq2^52^.

### Single cell analysis

Normal bone marrow(nBM) cells from Abbas et al.^47^ were processed using the standard *Seurat*^62^ workflow. Briefly data was normalized and scaled using the functions *NormalizeData()* with *scale.factor = 10,000* and *ScaleData()* respectively. 2000 variable genes were identified using *FindVariableFeatures()*. *RunPCA(), FindNeighbors(),FindClusters()* and *RunUMAP()* were run to cluster the data and generate a uMAP embedding using 30 PCs selected by inspecting the elbow plots. *FindAllMarkers()* was used to test differential expression of pro-survival (*BCL2*, *BCL2L2*, *BCL2L1*, *BCL2A1*, and *MCL1*) genes, in more differentiated cell-types relative to HSC and Progenitor cells, with *logfc.threshold* = 0 and *min.pct* = 0.05 using *wilcox test*. DEGs were identified at adjusted_pvalue < 0.01 and absolute average log2FC > 0.25.

Malignant cells from samples in Van Galen et al.^32^ (SeqWell) and Kuusanmaki et al. ^20^ (10x genomics), as defined in the original studies, were selected, and merged. The merged Seurat object was then processed, briefly data was normalized and scaled using the functions *NormalizeData()* with *scale.factor = 10,000* and *ScaleData()* respectively. 2000 variable genes were identified using *FindVariableFeatures()*. PCA was performed *RunPCA()* with the *npcs* = 100. The samples were integrated with Harmony^69^ using the *RunHarmony()* function, with the first 75 PCs while controlling for sample membership and sequencing technology. uMAP embedding was generated using *RunUMAP()* with *dims* = 75 and *reduction = “harmony*”.

Single cell gene expression data from pre- and post-treatment cells from a patient treated with VEN+AZA from Pie et al.^14^ was processed as described above for nBM. *FindTransferAnchors* and *TransferData* were used to transfer cell type labels from normal BM cells (**Figure 6A**). Pathway activity of Hallmark pathways in each cell was quantified using *VAM*^70^. Differentially active pathways were identified using the functions *FindAllMarkers()*, significant pathways were identified at absolute log 2FC > 0.15 and adjusted p-value < 0.1.

### Statistical analysis

Statistical analysis was performed in R (v 4.0.1) and GraphPad (v9). Association between continuous variables was quantified using spearman correlation. Differences between continuous variables was test using 1. Two-sided Wilcox test/T-test as indicated in case of two groups. 2. ANOVA was used in case of more than two groups and post-hoc pairwise testing was performed with Tukey’s test. Association between factor variables was performed using the Chi-square test. Correction for multiple testing was performed with FDR using the function *p.adjust*(). Significance was defined a p-value < 0.05 or q/adjusted p-value < 0.1 in case of multiple testing, unless specified.

## Acknowledgments

This work was supported by the National Cancer Institute’s Office of Cancer Genomics Cancer Target Discovery and Development (CTD^2) initiative. This work was supported in part by NIH (U01 CA247760) and the CPRIT (RP180248) to K.C., NCI (U01 CA217842) to G.B.M. and the NIH/NCI (R01CA235622) to M.K.. The results published here are based in part upon data generated by CTD^2 Network (https://www.cancer.gov/ccg/research/functional-genomics/ctd2/data-portal) established by the National Cancer Institute’s Office of Cancer Genomics.

## Author Contribution

V.M. and K.C. designed the study. V.M., K.C. and M.K. supervised the study. V.M., N.B., Y.H., C.L.R., L.M.C., S.H., R.I. performed the analysis. V.M., K.C., N.B., Y.H., C.L.R., L.M.C., S.H., R.I., M.D., J.W.T., G.B.M. and M.K. helped analyze the results. All authors helped to write the manuscript. All authors have read and approved this paper.

## Supplementary Figure legends

**Figure S1: NMF decomposition: A)** Heatmap of AUC percentiles for cell-lines across tissues of origin in CCLE. **B)** Boxplot of VEN AUC in resistant and sensitive cell-lines (two-sided T-test). **C)** Pathways enriched (p< 0.05 and q < 0.1) in the VEN resistance signature before and after filtering genes (see **Methods**). **D)** Plot of change in RSS for each increase in number of components used for the NMF. **E)** Heatmap of spearman correlation between patient component loadings. Correlation coefficients are reported in the cells. **F)** Heatmap of spearman correlation of patient level NMF component loadings with pathway activity of hallmark pathways. Digits in each cell are q-values after FDR correction across pathways for each component (column).

**Figure S2: Phenotypic, genetic, and transcriptional characteristics of VRS (in BeatAML1): A)** Mutually exclusive mutations in VR_C2 (top) and VR_C3 (bottom; see **Methods**). **B)** Heatmap of spearman correlation between VEN AUC and AML blast-type abundance scores. q-values after FDR correction are reported in the cells. **C)** Boxplot of AML blast-types with significantly different abundances (q < 0.1) between VEN resistant and sensitive samples (two-sided Wilcox-test followed by FDR correction is used to compute q-values.). **D)** Distribution of AML blast-type abundances in each VRS. **E)** Heatmap of pairwise difference between means abundance of AML blast-types in **Figure 2D** that showed significant differences between VRS. The pairwise difference in means is reported in cases where the difference is significant (Tukey’s test p.adj < 0.1)**: F)** Differentially activity (q < 0.1) pathways in VEN resistant patients relative to sensitive identified using GSEA. Red: over-expression and blue: suppression. **G)** Heatmap of activity of signaling pathways inferred using *Progeny* across patients in BeatAML1. **H)** Heatmap of T-statistics comparing activity of progeny pathways in VR_Cs vs VSC (left) and each VR_C vs other VR_Cs (right). P-values are computed with two-sided T-test, q-values after FDR correction for each column are reported in the cells. Significant at q < 0.1. **I)** Heatmap of TF-activity across patients that are differentially active (see **Methods**) in VR_C2 relative to other VRS. **J)** Heatmap of log2FC of *HOXA* and *HOXB* genes across comparisons, log2FC are reported in cases where the p.adj < 0.1. **K)** Heatmap of log2FC of genes for cytotoxicity (*PRF1*, *GZMA* and *GZMB*) and targets of immune checkpoint therapies (*PDCD1* and *CTLA4*), log2FC are reported in cases where the p.adj < 0.1. ***L)*** Boxplot of CTL abundance scores across VRS. P-values is computed with ANOVA and pairwise p.adj with a Tukey’s test. Heatmap of Z-transformed average AUC values of drugs that show specific sensitivity in **M)** VEN resistant samples relative to VSC and **N)** in each VR_C relative to other samples (see **Methods** for details). In **N,** average AUC of a drug is reported when it is specifically sensitive in a VR_C.

**Figure S3: Projecting VRS definitions onto new samples: A)** Schema to project VRS definitions onto target gene expression data (GE target) based on gene PCA loadings (gPC) computed from the BeatAML1 gene expression data (GE BeatAML_D) used in the NMF (**Figure 1**). Transpose of GE Target is multiplied with gPC to get PCA components of the target samples (tPC) thus projecting the samples into the same PCA space as BeatAML1 samples. Cluster definitions are assigned to the new samples (in GE Target) using k-means clustering based on centroid of clusters in BeatAML1 (see **Methods** for details). Heatmap of fraction of projected VRS definitions that are resistant and sensitive to VEN for **B)** BeatAML2 and **C)** CCLE. Association between VEN resistance status and VRS definitions is tested using Chi-square test. Boxplot comparing VEN AUC values between projected VRS definitions in **D)** BeatAML2 and **E)** CCLE. ANOVA is used to compute significance of difference between groups and pairwise tests comparing VR_Cs to VSC **(D)** and VSC+VR_C4 **(E)** was performed using Tukey’s test and significance defined at p.adj < 0.1. **Note:** VSC and VR_C4 are merged in CCLE cell-lines because of the small number of cell-lines and the transcriptional similarity between VSC and VR_C4 (**Figure 2E-F**). Heatmap of fractions of samples carrying a mutation in a gene in each VRS for **F)** BeatAML2 and **G)** TCGA. Only genes which show significant association with VRS states (q < 0.1; Chi-square test followed by FDR correction) were plotted, and testing was limited to mutations identified in BeatAML1 (**Figure 2C**, see **Methods**). **H)** Fractions of VR_C specific drugs discovered in BeatAML1 recaptured based on VRS definitions projected onto samples from BeatAML2 (see **Methods** for details). Same as **Figure S2E** for **I)** BeatAML2 and **J)** TCGA.

**Figure S4: Transcriptional characteristics of VRS projected onto new samples:** Same as **Figure 2 E-F** for VRS definitions projected onto **A)** BeatAML2 and **B)** TCGA samples. **C)** Same as **Figure S2L** for BeatAML2 (top) and TCGA (bottom). **D)** Same as **Figure S2K** for BeatAML2 (top) and TCGA (bottom). **E)** Same as **Figure S2J** for BeatAML2 (top) and TCGA (bottom).

**Figure S5: Developmental patterns associated with VRS:** Heatmap of fraction of samples in each VRS across FAB classifications in **A)** TCGA and **B)** CCLE. The significance of association between FAB and VRS membership is tested using the Chi-square test. **B)** Heatmap of log2FC of hematopoietic cell type markers in each VRS relative to all other VRS for BeatAML1 (left), BeatAML2 (mid) and TCGA (right). Log2FC are reported in the cells where p.adj < 0.1. **C)** PCA plot based on top 5000 variably expressed genes in sorted populations of AML blasts. Samples are colored by clinical classification of the sample and shapes indicate its cellular phenotype. **Note:** samples segregate by cell-type but not clinical classification. **D)** uMAP of healthy BM cells colored by cell-type.

**Figure S6: VR_C3 is characterized by activation of JAK-STAT signaling and is sensitive to its inhibition: A)** KEGG network for JAK-STAT signaling pathway colored by log2FC of genes when VR_C3 is compared to all other VRS in BeatAML1. **B)** Same as **Figure 4B** for BeatAML2 (top) and TCGA (bottom). **C)** Same as **Figure 4C** for BeatAML2 (top) and TCGA (bottom). **D)** Boxplot comparing drug AUC, corresponding to **Figure 4F**, in VR_C3 and non-VR_C3 cell-lines. P-value computed using two-sided T-test. **E)** Same as **Figure 4G** for HEL (top) and OCIM1 (bottom). **F)** Drug synergy plots for VEN and RUXO in F36P (left) and TF1(right; see **Methods**).

**Figure S7: VR_C1 is characterized by a transcriptional signature indicating active metabolism correlated with high PI3K signaling: A)** Heatmap of log2FC of metabolic enzymes and genes (in glycolysis, TCA cycle, glutaminolysis and ETC) across comparisons in BeatAML1 (left), BeatAML2 (mid) and TCGA (right). Log2FC are reported for genes with p.adj < 0.1. **B)** Heatmap of difference in means activity of nutrient transport pathways in VR_C1 relative to other VRS in BeatAML1 (left), BeatAML2 (mid) and TCGA (right) for nutrient transporter pathways with significant difference in activity across VRS (q < 0.1, ANOVA followed by FDR correction). Significance of pairwise differences was tested using Tukey’s test, and the difference is reported in the cells when significant (p.adj < 0.1). **C)** Same as **Figure 5B** for BeatAML2 (left) and TCGA (right). **D)** Same as **Figure 5C** for BeatAML2.

**Figure S8: VR_C1 is sensitive to Rapamycin, Panobinostat and CDK inhibition: A)** Boxplot of AUC for rapamycin (left), panobinostat (mid) and SNS-032 (right) in BeatAML1 (top) and BeatAML2 (bottom). Drugs show specificity to VR_C1 (Table 2). In each dataset drug pair one-sided T-tests followed by FDR correction was used to test if AUC in VR_C1 is significantly (q < 0.1) lower (sensitive) relative to other VRS. **B)** Pathways significantly (q< 0.1) activated and suppressed in AML blasts treated with panobinostat compared to control identified using GSEA. positive stat.mean (mean statistic, see **Methods**) indicates over-expression and negative indicates suppression. **C)** heatmap of drug-response AUCs for cell-lines treated with VEN, PANO and VEN+PANO (left) and boxplots (right) comparing AUC between VR_C1-like cell-lines and other cell-lines (p-values computed using two-sided T-test). **D)** Boxplot of drug-response AUC for mTOR inhibitors. One-sided T-test followed by FDR correction was used to identify drugs with significantly lower AUC (q < 0.1) in VR_C1-like cell-lines. **E)** GSEA as in **Figure S8B** for SNS-032 treated MOLT4 cells relative to untreated cells. Pathways are plotted at a lowered threshold for significance (q < 0.2) with the q-values reported alongside the pathway labels.

