## Supplements for "Transcriptional and phenotypic heterogeneity underpinning venetoclax resistance in AML"

### SuppFig 1

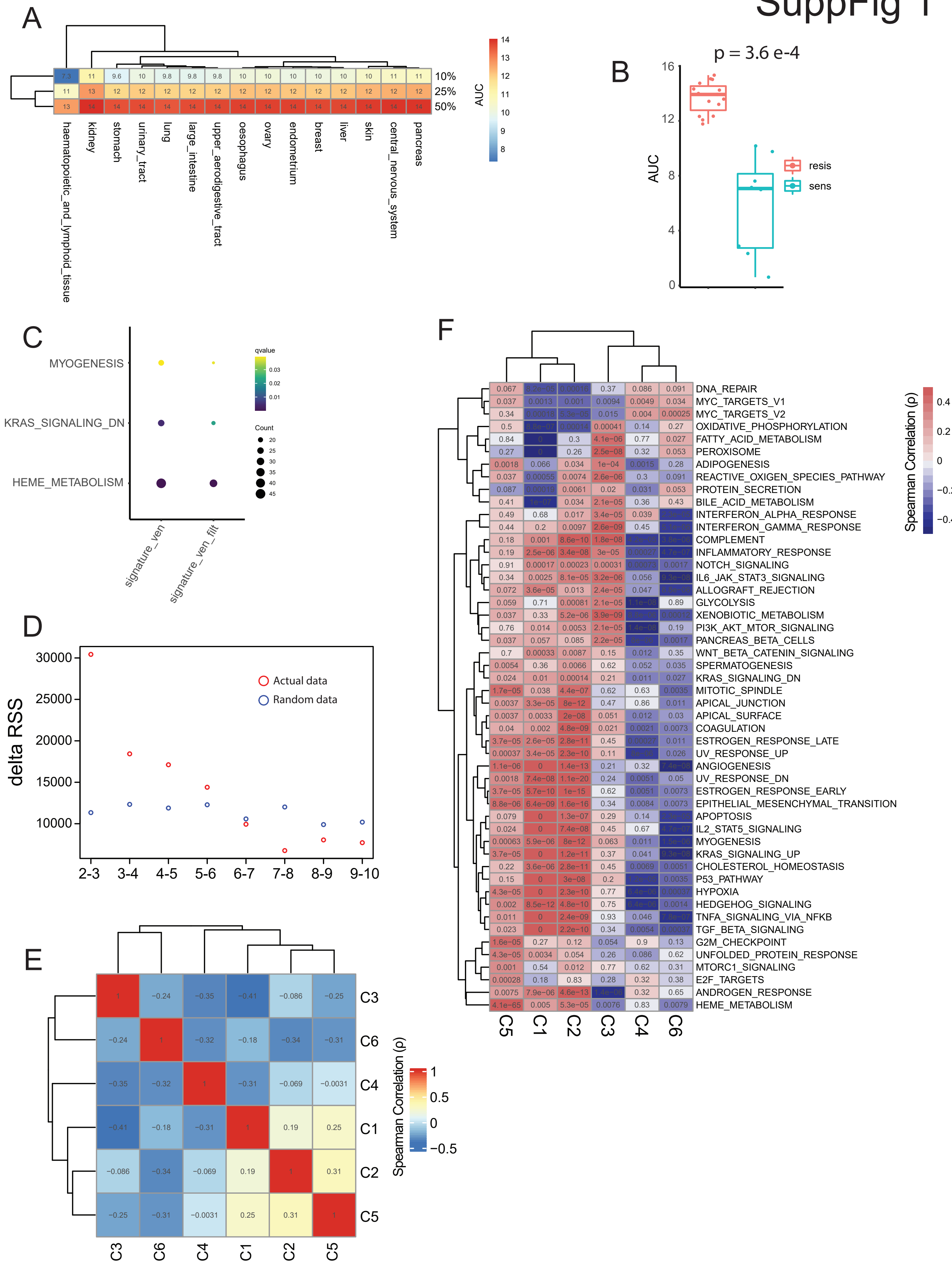
A
10%
25%
50%
AUC
haematopoietic_and_lymphoid_tissue
kidney
stomach
urinary_tract
lung
large_intestine
upper_aerodigestive_tract
oesophagus
ovary
endometrium
breast
liver
skin
central_nervous_system
pancreas
B
p = 3.6 e-4
AUC
resis
sens
C
MYOGENESIS
KRAS_SIGNALING_DN
HEME_METABOLISM
qvalue
Count
signature_ven
signature_ven_filt
D
delta RSS
Actual data
Random data
2-3
3-4
4-5
5-6
6-7
7-8
8-9
9-10
E
C3
C6
C4
C1
C2
C5
Spearman Correlation (ρ)
F
DNA_REPAIR
MYC_TARGETS_V1
MYC_TARGETS_V2
OXIDATIVE_PHOSPHORYLATION
FATTY_ACID_METABOLISM
PEROXISOME
ADIPOGENESIS
REACTIVE_OXIGEN_SPECIES_PATHWAY
PROTEIN_SECRETION
BILE_ACID_METABOLISM
INTERFERON_ALPHA_RESPONSE
INTERFERON_GAMMA_RESPONSE
COMPLEMENT
INFLAMMATORY_RESPONSE
NOTCH_SIGNALING
IL6_JAK_STAT3_SIGNALING
ALLOGRAFT_REJECTION
GLYCOLYSIS
XENOBIOTIC_METABOLISM
PI3K_AKT_MTOR_SIGNALING
PANCREAS_BETA_CELLS
WNT_BETA_CATENIN_SIGNALING
SPERMATOGENESIS
KRAS_SIGNALING_DN
MITOTIC_SPINDLE
APICAL_JUNCTION
APICAL_SURFACE
COAGULATION
ESTROGEN_RESPONSE_LATE
UV_RESPONSE_UP
ANGIOGENESIS
UV_RESPONSE_DN
ESTROGEN_RESPONSE_EARLY
EPITHELIAL_MESENCHYMAL_TRANSITION
APOPTOSIS
IL2_STAT5_SIGNALING
MYOGENESIS
KRAS_SIGNALING_UP
CHOLESTEROL_HOMEOSTASIS
P53_PATHWAY
HYPOXIA
HEDGEHOG_SIGNALING
TNFA_SIGNALING_VIA_NFKB
TGF_BETA_SIGNALING
G2M_CHECKPOINT
UNFOLDED_PROTEIN_RESPONSE
MTORC1_SIGNALING
E2F_TARGETS
ANDROGEN_RESPONSE
HEME_METABOLISM
C5
C1
C2
C3
C4
C6
Spearman Correlation (ρ)

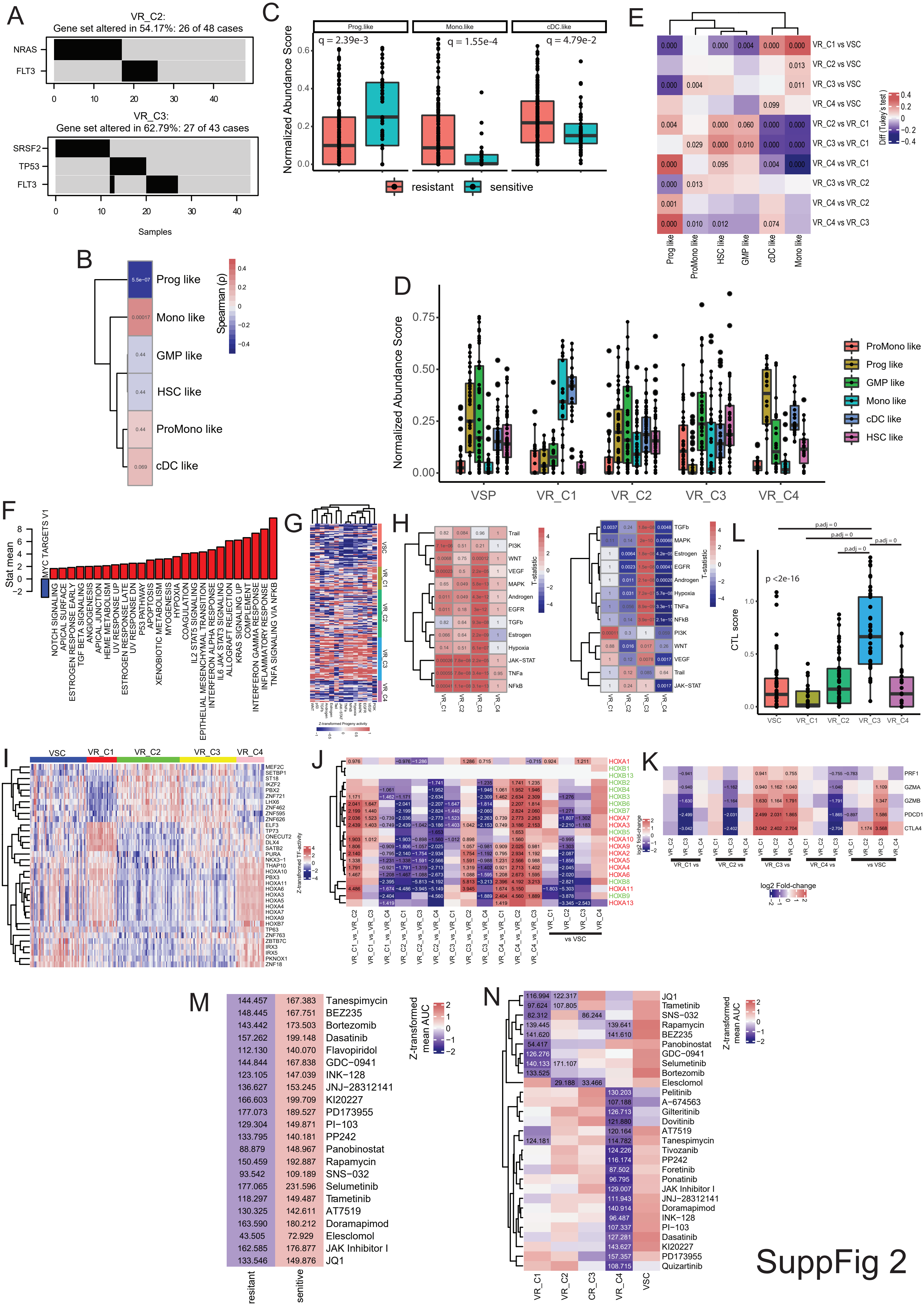

A
VR_C2:
Gene set altered in 54.17%: 26 of 48 cases
NRAS
FLT3
VR_C3:
Gene set altered in 62.79%: 27 of 43 cases
SRSF2
TP53
FLT3
Samples
B
Prog like
Mono like
GMP like
HSC like
ProMono like
cDC like
Spearman (ρ)
C
Prog.like
Mono.like
cDC.like
q = 2.39e-3
q = 1.55e-4
q = 4.79e-2
Normalized Abundance Score
resistant
sensitive
D
Normalized Abundance Score
VSP
VR_C1
VR_C2
VR_C3
VR_C4
ProMono like
Prog like
GMP like
Mono like
cDC like
HSC like
E
VR_C1 vs VSC
VR_C2 vs VSC
VR_C3 vs VSC
VR_C4 vs VSC
VR_C2 vs VR_C1
VR_C3 vs VR_C1
VR_C4 vs VR_C1
VR_C3 vs VR_C2
VR_C4 vs VR_C2
VR_C4 vs VR_C3
Diff (Tukey's test)
Prog like
ProMono like
HSC like
GMP like
cDC like
Mono like
F
Stat mean
MYC TARGETS V1
NOTCH SIGNALING
APICAL SURFACE
ESTROGEN RESPONSE EARLY
TGF BETA SIGNALING
ANGIOGENESIS
APICAL JUNCTION
HEME METABOLISM
UV RESPONSE UP
ESTROGEN RESPONSE LATE
UV RESPONSE DN
P53 PATHWAY
APOPTOSIS
XENOBIOTIC METABOLISM
MYOGENESIS
HYPOXIA
COAGULATION
IL2 STAT5 SIGNALING
EPITHELIAL MESENCHYMAL TRANSITION
INTERFERON ALPHA RESPONSE
IL6 JAK STAT3 SIGNALING
ALLOGRAFT REJECTION
KRAS SIGNALING UP
COMPLEMENT
INTERFERON GAMMA RESPONSE
INFLAMMATORY RESPONSE
TNFA SIGNALING VIA NFKB
G
VSC
VR_C1
VR_C2
VR_C3
VR_C4
Z-transformed Progeny activity
H
Trail
PI3K
WNT
VEGF
MAPK
Androgen
EGFR
TGFb
Estrogen
Hypoxia
JAK-STAT
TNFa
NFkB
T-statistic
L
p.adj = 0
p <2e-16
CTL score
VSC
VR_C1
VR_C2
VR_C3
VR_C4
I
VSC
VR_C1
VR_C2
VR_C3
VR_C4
Z-transformed TF-activity
J
HOXA1
HOXB1
HOXB13
HOXB2
HOXB4
HOXB3
HOXB6
HOXB7
HOXA7
HOXA3
HOXB5
HOXA10
HOXA9
HOXA2
HOXA5
HOXA4
HOXA6
HOXB8
HOXA11
HOXB9
HOXA13
log2 fold-change
vs VSC
K
PRF1
GZMA
GZMB
PDCD1
CTLA4
VR_C1 vs
VR_C2 vs
VR_C3 vs
VR_C4 vs
vs VSC
log2 Fold-change
M
Tanespimycin
BEZ235
Bortezomib
Dasatinib
Flavopiridol
GDC−0941
INK−128
JNJ−28312141
KI20227
PD173955
PI−103
PP242
Panobinostat
Rapamycin
SNS−032
Selumetinib
Trametinib
AT7519
Doramapimod
Elesclomol
JAK Inhibitor I
JQ1
resitant
senitive
Z-transformed mean AUC
N
JQ1
Trametinib
SNS−032
Rapamycin
BEZ235
Panobinostat
GDC−0941
Selumetinib
Bortezomib
Elesclomol
Pelitinib
A−674563
Gilteritinib
Dovitinib
AT7519
Tanespimycin
Tivozanib
PP242
Foretinib
Ponatinib
JAK Inhibitor I
JNJ−28312141
Doramapimod
INK−128
PI−103
Dasatinib
KI20227
PD173955
Quizartinib
VR_C1
VR_C2
CR_C3
VR_C4
VSC
Z-transformed mean AUC


SuppFig 2

SuppFig 3

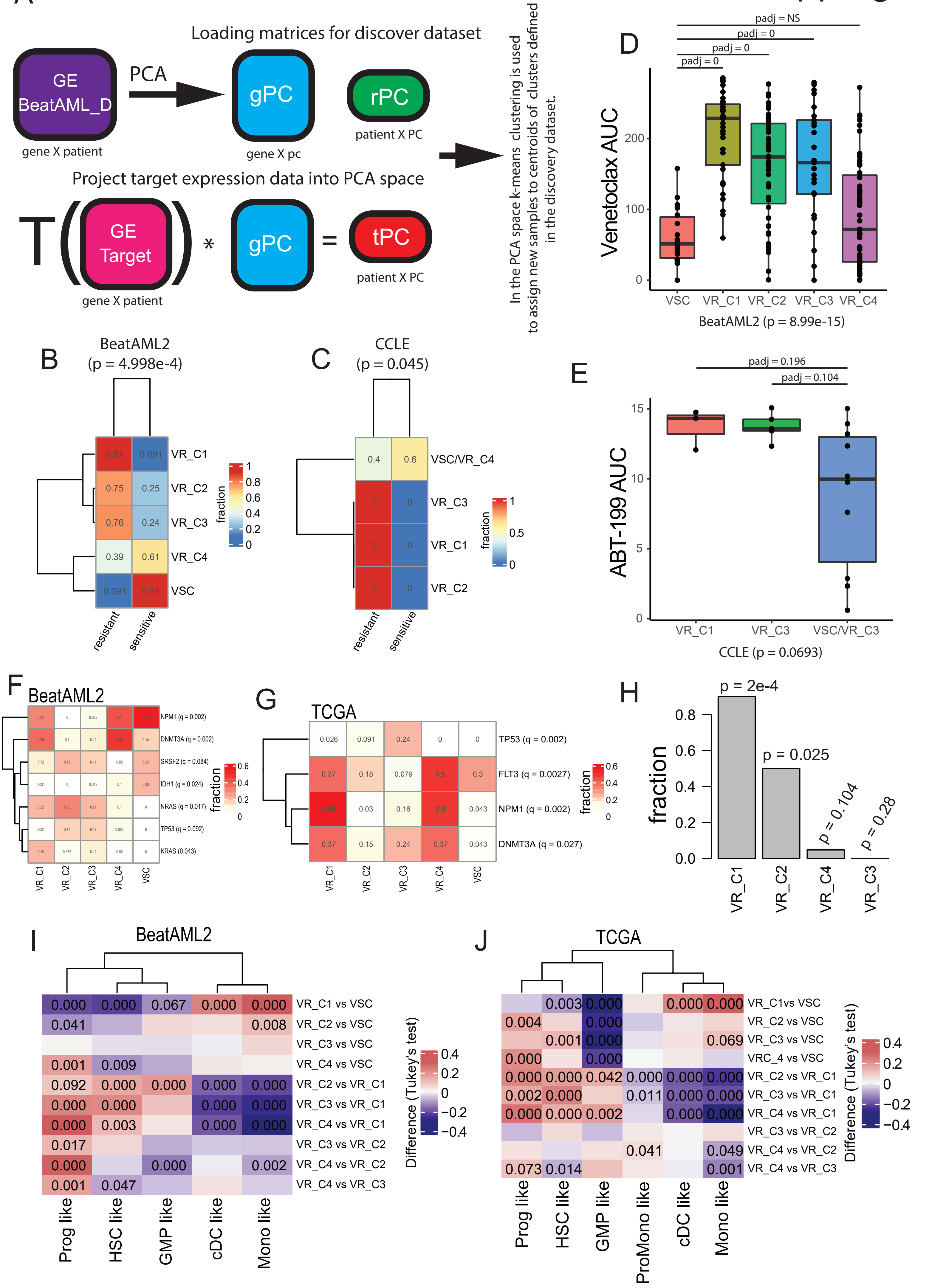
A
Loading matrices for discover dataset
GE
BeatAML_D
gene X patient
PCA
gPC
gene X pc
rPC
patient X PC
Project target expression data into PCA space
T
GE
Target
gene X patient
*
gPC
=
tPC
patient X PC
In the PCA space k-means clustering is used to assign new samples to centroids of clusters defined in the discovery dataset.
D
padj = NS
padj = 0
padj = 0
padj = 0
Venetoclax AUC
VSC
VR_C1
VR_C2
VR_C3
VR_C4
BeatAML2 (p = 8.99e-15)
B
BeatAML2
(p = 4.998e-4)
VR_C1
VR_C2
VR_C3
VR_C4
VSC
resistant
sensitive
fraction
C
CCLE
(p = 0.045)
VSC/VR_C4
VR_C3
VR_C1
VR_C2
resistant
sensitive
fraction
E
padj = 0.196
padj = 0.104
ABT-199 AUC
VR_C1
VR_C3
VSC/VR_C3
CCLE (p = 0.0693)
F
BeatAML2
NPM1 (q = 0.002)
DNMT3A (q = 0.002)
SRSF2 (q = 0.084)
IDH1 (q = 0.024)
NRAS (q = 0.017)
TP53 (q = 0.092)
KRAS (0.043)
fraction
VR_C1
VR_C2
VR_C3
VR_C4
VSC
G
TCGA
TP53 (q = 0.002)
FLT3 (q = 0.0027)
NPM1 (q = 0.002)
DNMT3A (q = 0.027)
fraction
VR_C1
VR_C2
VR_C3
VR_C4
VSC
H
p = 2e-4
p = 0.025
p = 0.104
p = 0.28
fraction
VR_C1
VR_C2
VR_C4
VR_C3
I
BeatAML2
VR_C1 vs VSC
VR_C2 vs VSC
VR_C3 vs VSC
VR_C4 vs VSC
VR_C2 vs VR_C1
VR_C3 vs VR_C1
VR_C4 vs VR_C1
VR_C3 vs VR_C2
VR_C4 vs VR_C2
VR_C4 vs VR_C3
Difference (Tukey's test)
Prog like
HSC like
GMP like
cDC like
Mono like
J
TCGA
VR_C1vs VSC
VR_C2 vs VSC
VR_C3 vs VSC
VRC_4 vs VSC
VR_C2 vs VR_C1
VR_C3 vs VR_C1
VR_C4 vs VR_C1
VR_C3 vs VR_C2
VR_C4 vs VR_C2
VR_C4 vs VR_C3
Difference (Tukey's test)
Prog like
HSC like
GMP like
ProMono like
cDC like
Mono like

### SuppFig 4

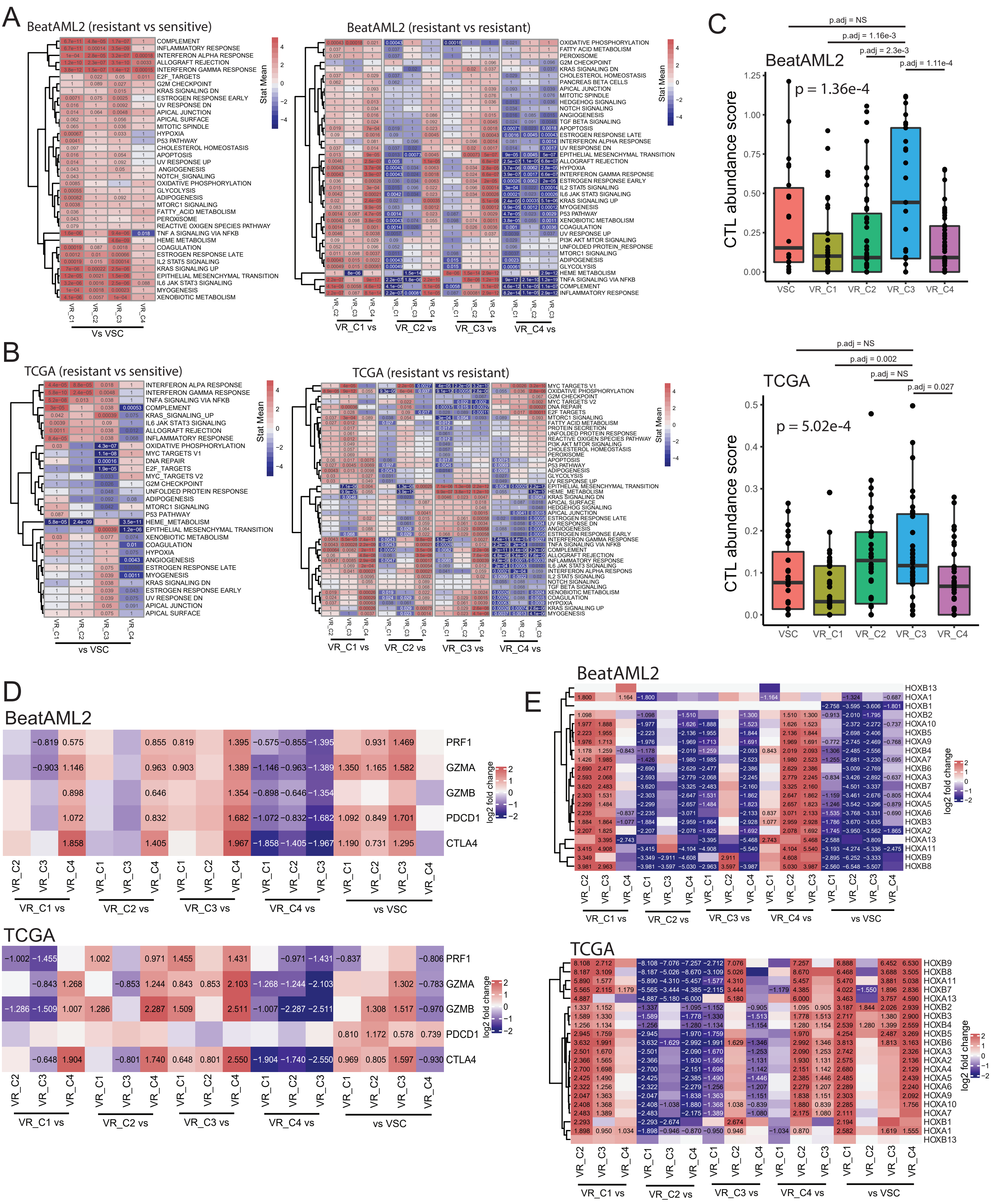
A
BeatAML2 (resistant vs sensitive)
BeatAML2 (resistant vs resistant)
Stat Mean
Vs VSC
VR_C1 vs
VR_C2 vs
VR_C3 vs
VR_C4 vs
B
TCGA (resistant vs sensitive)
TCGA (resistant vs resistant)
vs VSC
C
BeatAML2
p = 1.36e-4
p.adj = NS
p.adj = 1.16e-3
p.adj = 2.3e-3
p.adj = 1.11e-4
CTL abundance score
TCGA
p = 5.02e-4
p.adj = NS
p.adj = 0.002
p.adj = NS
p.adj = 0.027
VSC
VR_C1
VR_C2
VR_C3
VR_C4
D
BeatAML2
TCGA
PRF1
GZMA
GZMB
PDCD1
CTLA4
log2 fold change
E
BeatAML2
TCGA
vs VSC

### SuppFig 5

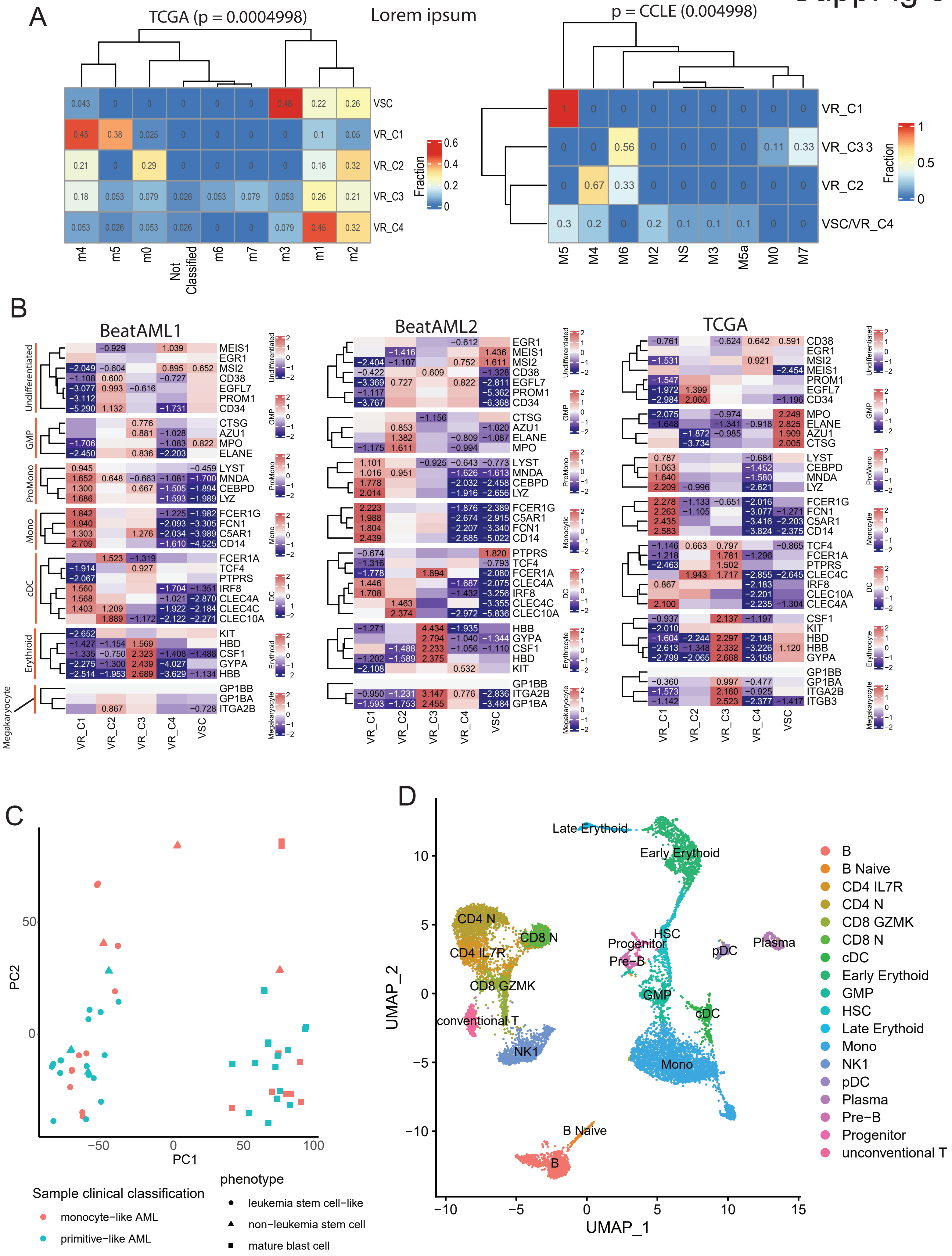

A
TCGA (p = 0.0004998)
Lorem ipsum
p = CCLE (0.004998)
VSC
VR_C1
VR_C2
VR_C3
VR_C4
m4
m5
m0
Not Classified
m6
m7
m3
m1
m2
Fraction
VR_C33
VSC/VR_C4
M5
M4
M6
M2
NS
M3
M5a
M0
M7
B
BeatAML1
BeatAML2
TCGA
Undifferentiated
GMP
ProMono
Mono
Monocytic
cDC
DC
Erythroid
Erythrocyte
Megakaryocyte
MEIS1
EGR1
MSI2
CD38
EGFL7
PROM1
CD34
CTSG
AZU1
MPO
ELANE
LYST
MNDA
CEBPD
LYZ
FCER1G
FCN1
C5AR1
CD14
FCER1A
TCF4
PTPRS
IRF8
CLEC4A
CLEC4C
CLEC10A
KIT
HBD
CSF1
GYPA
HBB
GP1BB
GP1BA
ITGA2B
ITGB3
VR_C1
VR_C2
VR_C3
VR_C4
VSC
C
PC2
PC1
Sample clinical classification
monocyte−like AML
primitive−like AML
phenotype
leukemia stem cell−like
non−leukemia stem cell
mature blast cell
D
UMAP_2
UMAP_1
Late Erythoid
Early Erythoid
CD4 N
CD8 N
CD4 IL7R
CD8 GZMK
conventional T
NK1
Progenitor
Pre−B
HSC
pDC
Plasma
GMP
cDC
Mono
B Naive
B
B Naive
CD4 IL7R
CD4 N
CD8 GZMK
CD8 N
cDC
Early Erythoid
GMP
HSC
Late Erythoid
Mono
NK1
pDC
Plasma
Pre−B
Progenitor
unconventional T

### SuppFig 6

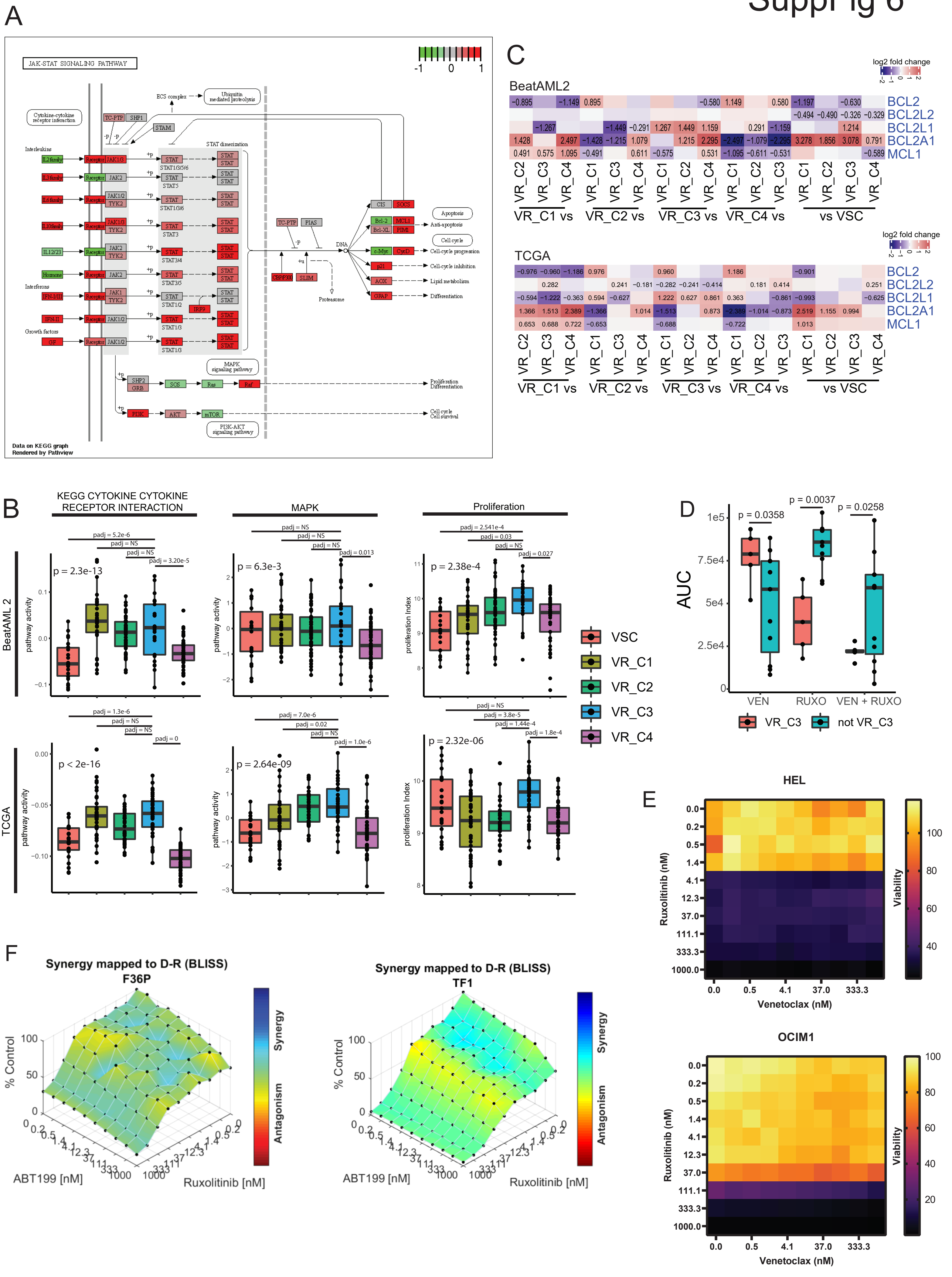
A
JAK-STAT SIGNALING PATHWAY
-1 0 1
Cytokine-cytokine receptor interaction
ECS complex
Ubiquitin mediated proteolysis
Interleukins
Interferons
Growth factors
STAT dimerization
TC-PTP SHP1
STAM
CIS SOCS
Bcl-2 MCL1
Bcl-XL PIM1
c-Myc CycD
p21
AOX
GFAP
Apoptosis
Anti-apoptosis
Cell cycle
Cell-cycle progression
Cell-cycle inhibition
Lipid metabolism
Differentiation
DNA
PIAS
CBP/P300 SLIM
Proteasome
MAPK signaling pathway
SHP2 GRB SOS Ras Raf
PI3K AKT mTOR
PI3K-AKT signaling pathway
Proliferation Differentiation
Cell cycle Cell survival
Data on KEGG graph
Rendered by Pathview
B
KEGG CYTOKINE CYTOKINE RECEPTOR INTERACTION
MAPK
Proliferation
BeatAML 2
TCGA
pathway activity
proliferation Index
p = 2.3e-13
p = 6.3e-3
p = 2.38e-4
p < 2e-16
p = 2.64e-09
p = 2.32e-06
padj = 5.2e-6
padj = NS
padj = 3.20e-5
padj = 0.013
padj = 2.541e-4
padj = 0.03
padj = 0.027
padj = 1.3e-6
padj = 0
padj = 7.0e-6
padj = 0.02
padj = 1.0e-6
padj = 3.8e-5
padj = 1.44e-4
padj = 1.8e-4
VSC
VR_C1
VR_C2
VR_C3
VR_C4
C
BeatAML2
TCGA
log2 fold change
BCL2
BCL2L2
BCL2L1
BCL2A1
MCL1
VR_C1 vs
VR_C2 vs
VR_C3 vs
VR_C4 vs
vs VSC
D
AUC
p = 0.0358
p = 0.0037
p = 0.0258
VEN
RUXO
VEN + RUXO
VR_C3
not VR_C3
E
HEL
OCIM1
Ruxolitinib (nM)
Venetoclax (nM)
Viability
F
Synergy mapped to D-R (BLISS)
F36P
TF1
% Control
ABT199 [nM]
Ruxolitinib [nM]
Synergy
Antagonism

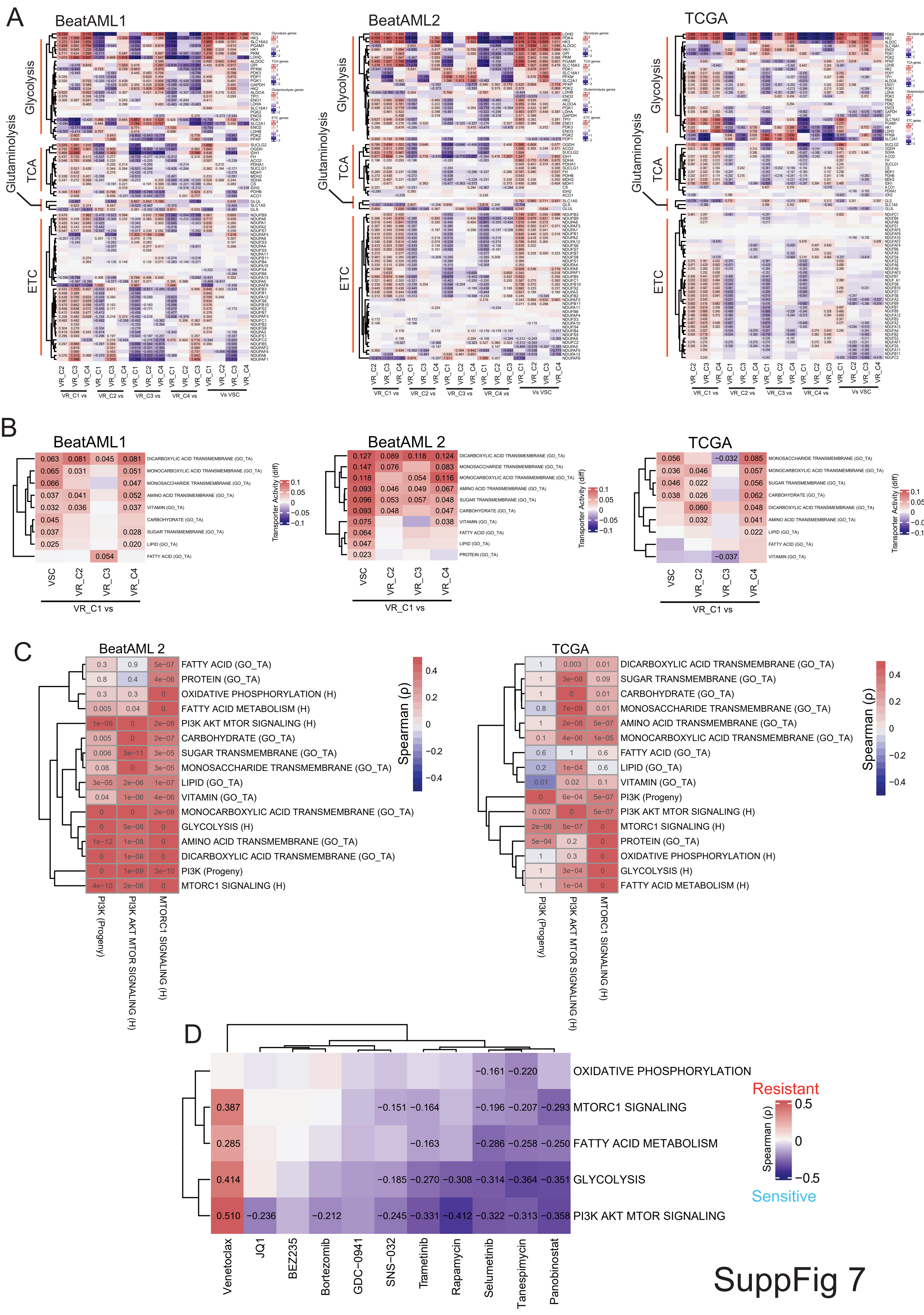
A
BeatAML1
BeatAML2
TCGA
Glycolysis
TCA
Glutaminolysis
ETC
VR_C1 vs
VR_C2 vs
VR_C3 vs
VR_C4 vs
Vs VSC
B
BeatAML1
0.063 0.081 0.045 0.081 DICARBOXYLIC ACID TRANSMEMBRANE (GO_TA)
0.065 0.031 0.051 MONOCARBOXYLIC ACID TRANSMEMBRANE (GO_TA)
0.066 0.047 MONOSACCHARIDE TRANSMEMBRANE (GO_TA)
0.037 0.041 0.052 AMINO ACID TRANSMEMBRANE (GO_TA)
0.032 0.036 0.037 VITAMIN (GO_TA)
0.045 CARBOHYDRATE (GO_TA)
0.037 0.028 SUGAR TRANSMEMBRANE (GO_TA)
0.025 0.020 LIPID (GO_TA)
0.054 FATTY ACID (GO_TA)
VSC
VR_C2
VR_C3
VR_C4
VR_C1 vs
Transporter Activity (diff)
BeatAML 2
0.127 0.089 0.118 0.124 DICARBOXYLIC ACID TRANSMEMBRANE (GO_TA)
0.147 0.076 0.083 MONOSACCHARIDE TRANSMEMBRANE (GO_TA)
0.118 0.054 0.116 MONOCARBOXYLIC ACID TRANSMEMBRANE (GO_TA)
0.093 0.046 0.049 0.067 AMINO ACID TRANSMEMBRANE (GO_TA)
0.096 0.053 0.057 0.048 SUGAR TRANSMEMBRANE (GO_TA)
0.093 0.048 0.047 CARBOHYDRATE (GO_TA)
0.075 0.038 VITAMIN (GO_TA)
0.064 FATTY ACID (GO_TA)
0.047 LIPID (GO_TA)
0.023 PROTEIN (GO_TA)
TCGA
0.056 −0.032 0.085 MONOSACCHARIDE TRANSMEMBRANE (GO_TA)
0.036 0.046 0.057 MONOCARBOXYLIC ACID TRANSMEMBRANE (GO_TA)
0.046 0.022 0.056 SUGAR TRANSMEMBRANE (GO_TA)
0.038 0.026 0.062 CARBOHYDRATE (GO_TA)
0.060 0.048 DICARBOXYLIC ACID TRANSMEMBRANE (GO_TA)
0.032 0.041 AMINO ACID TRANSMEMBRANE (GO_TA)
0.022 LIPID (GO_TA)
FATTY ACID (GO_TA)
−0.037 VITAMIN (GO_TA)
C
BeatAML 2
0.3 0.9 5e-07 FATTY ACID (GO_TA)
0.8 0.4 4e-06 PROTEIN (GO_TA)
0.3 0.3 0 OXIDATIVE PHOSPHORYLATION (H)
0.005 0.04 0 FATTY ACID METABOLISM (H)
1e-09 0 2e-08 PI3K AKT MTOR SIGNALING (H)
0.005 0 2e-07 CARBOHYDRATE (GO_TA)
0.006 3e-11 3e-05 SUGAR TRANSMEMBRANE (GO_TA)
0.08 0 3e-05 MONOSACCHARIDE TRANSMEMBRANE (GO_TA)
3e-05 2e-06 1e-07 LIPID (GO_TA)
0.04 1e-06 4e-06 VITAMIN (GO_TA)
0 0 2e-08 MONOCARBOXYLIC ACID TRANSMEMBRANE (GO_TA)
0 5e-08 0 GLYCOLYSIS (H)
1e-12 1e-08 0 AMINO ACID TRANSMEMBRANE (GO_TA)
0 1e-08 0 DICARBOXYLIC ACID TRANSMEMBRANE (GO_TA)
0 1e-09 3e-10 PI3K (Progeny)
4e-10 2e-08 0 MTORC1 SIGNALING (H)
PI3K (Progeny)
PI3K AKT MTOR SIGNALING (H)
MTORC1 SIGNALING (H)
Spearman (ρ)
TCGA
1 0.003 0.01 DICARBOXYLIC ACID TRANSMEMBRANE (GO_TA)
1 3e-08 0.09 SUGAR TRANSMEMBRANE (GO_TA)
1 0 0.01 CARBOHYDRATE (GO_TA)
0.8 7e-09 0.01 MONOSACCHARIDE TRANSMEMBRANE (GO_TA)
1 2e-08 5e-07 AMINO ACID TRANSMEMBRANE (GO_TA)
0.1 4e-06 1e-05 MONOCARBOXYLIC ACID TRANSMEMBRANE (GO_TA)
0.6 1 0.6 FATTY ACID (GO_TA)
0.2 1e-04 0.6 LIPID (GO_TA)
0.01 0.02 0.1 VITAMIN (GO_TA)
0 6e-04 5e-07 PI3K (Progeny)
0.002 0 5e-07 PI3K AKT MTOR SIGNALING (H)
2e-06 5e-07 0 MTORC1 SIGNALING (H)
5e-04 0.2 0 PROTEIN (GO_TA)
1 0.3 0 OXIDATIVE PHOSPHORYLATION (H)
1 3e-04 0 GLYCOLYSIS (H)
1 1e-04 0 FATTY ACID METABOLISM (H)
D
−0.161 −0.220 OXIDATIVE PHOSPHORYLATION
0.387 −0.151 −0.164 −0.196 −0.207 −0.293 MTORC1 SIGNALING
0.285 −0.163 −0.286 −0.258 −0.250 FATTY ACID METABOLISM
0.414 −0.185 −0.270 −0.308 −0.314 −0.364 −0.351 GLYCOLYSIS
0.510 −0.236 −0.212 −0.245 −0.331 −0.412 −0.322 −0.313 −0.358 PI3K AKT MTOR SIGNALING
Venetoclax
JQ1
BEZ235
Bortezomib
GDC−0941
SNS−032
Trametinib
Rapamycin
Selumetinib
Tanespimycin
Panobinostat
Resistant
Sensitive
Spearman (ρ)

SuppFig 7

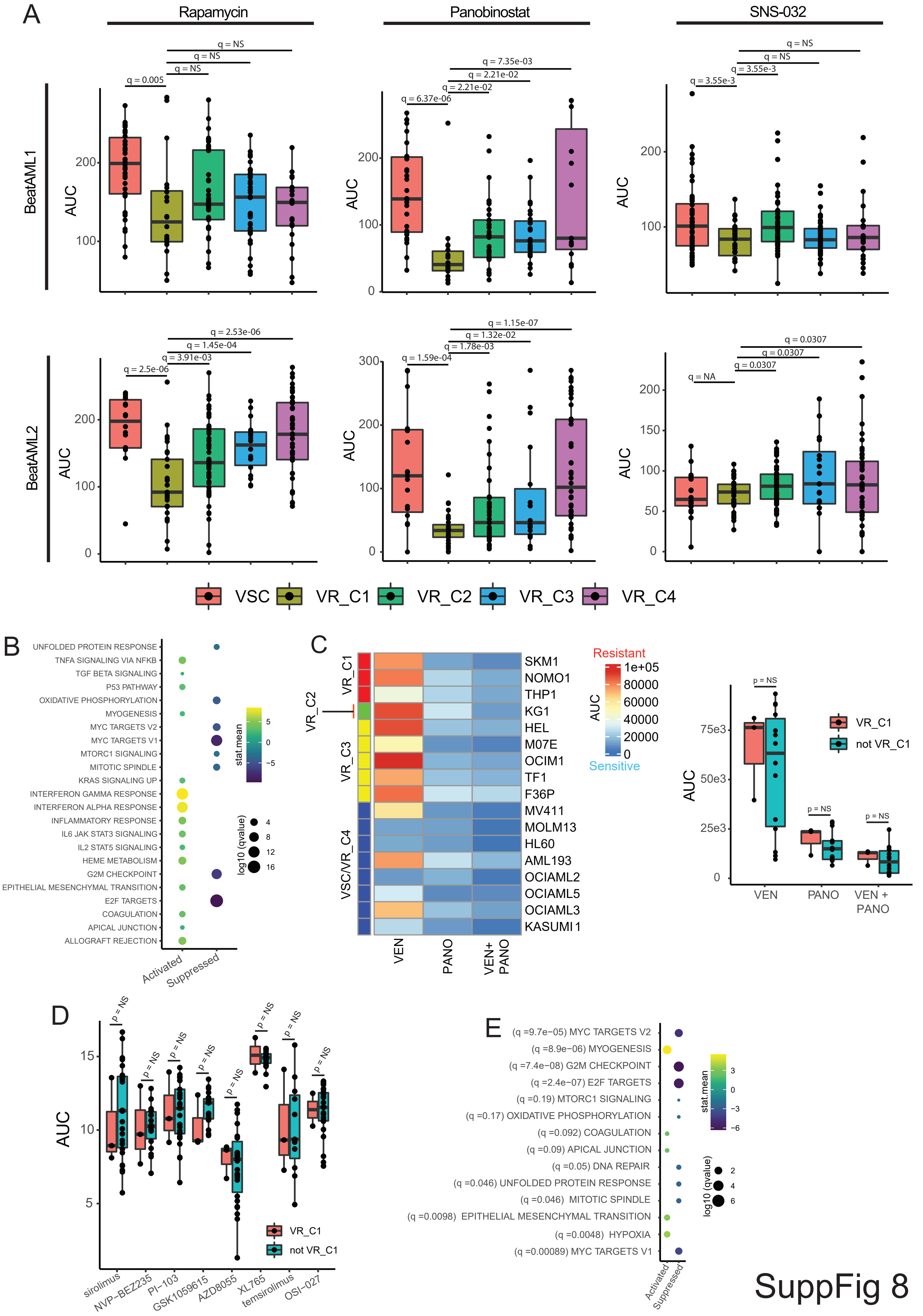
A
Rapamycin
Panobinostat
SNS-032
BeatAML1
BeatAML2
AUC
q = 0.005
q = NS
q = 6.37e-06
q = 2.21e-02
q = 7.35e-03
q = 3.55e-3
q = 2.5e-06
q = 3.91e-03
q = 1.45e-04
q = 2.53e-06
q = 1.59e-04
q = 1.78e-03
q = 1.32e-02
q = 1.15e-07
q = NA
q = 0.0307
VSC
VR_C1
VR_C2
VR_C3
VR_C4
B
UNFOLDED PROTEIN RESPONSE
TNFA SIGNALING VIA NFKB
TGF BETA SIGNALING
P53 PATHWAY
OXIDATIVE PHOSPHORYLATION
MYOGENESIS
MYC TARGETS V2
MYC TARGETS V1
MTORC1 SIGNALING
MITOTIC SPINDLE
KRAS SIGNALING UP
INTERFERON GAMMA RESPONSE
INTERFERON ALPHA RESPONSE
INFLAMMATORY RESPONSE
IL6 JAK STAT3 SIGNALING
IL2 STAT5 SIGNALING
HEME METABOLISM
G2M CHECKPOINT
EPITHELIAL MESENCHYMAL TRANSITION
E2F TARGETS
COAGULATION
APICAL JUNCTION
ALLOGRAFT REJECTION
Activated
Suppressed
stat.mean
log10 (qvalue)
C
VR_C1
VR_C2
VR_C3
VSC/VR_C4
SKM1
NOMO1
THP1
KG1
HEL
M07E
OCIM1
TF1
F36P
MV411
MOLM13
HL60
AML193
OCIAML2
OCIAML5
OCIAML3
KASUMI 1
VEN
PANO
VEN+ PANO
Resistant
AUC
1e+05
80000
60000
40000
20000
0
Sensitive
p = NS
VR_C1
not VR_C1
75e3
50e3
25e3
VEN
PANO
VEN + PANO
D
AUC
sirolimus
NVP-BEZ235
PI-103
GSK1059615
AZD8055
XL765
temsirolimus
OSI-027
E
(q =9.7e−05) MYC TARGETS V2
(q =8.9e−06) MYOGENESIS
(q =7.4e−08) G2M CHECKPOINT
(q =2.4e−07) E2F TARGETS
(q =0.19) MTORC1 SIGNALING
(q =0.17) OXIDATIVE PHOSPHORYLATION
(q =0.092) COAGULATION
(q =0.09) APICAL JUNCTION
(q =0.05) DNA REPAIR
(q =0.046) UNFOLDED PROTEIN RESPONSE
(q =0.046) MITOTIC SPINDLE
(q =0.0098) EPITHELIAL MESENCHYMAL TRANSITION
(q =0.0048) HYPOXIA
(q =0.00089) MYC TARGETS V1
stat.mean
log10 (qvalue)
Activated
Suppressed
SuppFig 8

**Supplementary Notes 1: Characteristics of VRS identified in BeatAML1 are conserved in VRS definitions projected onto samples from other datasets:**

VEN resistance status match projected VRS states in BeatAML2 and CCLE cell lines

VRS states projected onto BeatAML2 samples (**Figure S3B**) and CCLE cell lines (**Figure S3C**) show a strong association with VEN resistance status defined based on the AUC of VEN in the datasets (see **Methods**). In case of BeatAML2 (**Figure S3B**) VR_C1-3 show a strong enrichment (>=75%) for resistant samples and VSC shows an enrichment for sensitive samples (~91%). VR_C4 shows a mixture of resistant (39%) and sensitive samples (61%). The transcriptional similarity between VR_C4 and VSC (**Figure 2E**) likely makes the projection noisy. We therefore merge VR_C4 and VSC in the case of CCLE given the limited number of cell lines in the analysis (n = 37). In CCLE all VR_C clusters are exclusively VEN resistant cell-lines with VSC/VR_C4 showing an enrichment for sensitive cell-lines (60%; **Figure S3C**). Consistent with these observations VR_Cs show higher VEN AUC relative to VSC samples and significantly higher in the cases of VR_C1-3 in BeatAML2(**Figure S3D**). In CCLE for VR_Cs with more than one cell line where VEN response was measured (VR_C1 and 3), showed a trend towards higher VEN AUC relative to VSC/VR_C4, but the differences are only nominally significant (adjusted P-values of 0.196 and 0.104 for VR_C1 and 3 respectively; **Figure S3D**).

Conserved mutational patterns

To test conservation of mutational patterns we tested for association between projected VRS status and mutations in targets datasets (BeatAML2 and TCGA) as in the discovery datasets (BeatAML1; see **Methods**) but limited the testing to genes identified in the discovery cohort (**Figure 2C**). In BeatAML2 (**Figure S3F**) we re-captured high mutation rate of *NPM1* in VSC, VR_C1 and VR_C4. *DNMT3A* mutations were also more prevalent in VR_C1 and VR_C4. IDH1 mutations were most frequently observed in VSC (23%) unlike in BeatAML1, though we also saw modest enrichment (10%) in VR_C4 (**Figure 2C** and **Figure S3F**). RAS mutations were enriched in VR_C1 and 2, with VR_C2 showing a preference for *NRAS* mutations, also consistent with observations in BeatAML1 (**Figure 2C**). *TP53* and *SRSF2* mutations were more frequent in VR_C3 and VR_C2 as in the BeatAML1, however SRSF2 was also frequently found mutated in VSC which was not observed in the BeatAML1 (**Figure S3F** and **2C**). While fewer mutations were recapture in TCGA (4/8) we did recapture higher mutation rates of TP53 in VRC_3, higher rates of mutations of *FLT3* in VR_C1,4 and VSC and the higher mutations rates of *NMP1* and *DNMT3A* in VR_C1 and 4, though the high rates of *NMP1* in VSC (**Figure 2C**) was missing (**Figure S3G**). These data indicates that the genetic associations with VRS were largely preserved when VRS were projected onto target datasets (**Figure S3G-C** and **2C**).

Cluster specificity of individual drugs is conserved in BeatAML2

We next tested the sensitivity of cluster specific drugs identified in BeatAML1, utilizing the same procedure (see **Methods**; **Table 2**) in BeatAML2. We used a randomization approach to calculate if the rate at which cluster specific drugs were re-captured in each cluster were significant (see **Methods; Figure S3H**). The recapture rates for VR_C1 (90%) and VR_C2 (50%) were significant ($p < 0.05$), while the rate of recapture for VR_C4 was nominally significant (4.7% at $p = 0.104$). We recovered neither of the 2 drugs identified in case of VR_C3. Our VRS projections were therefore largely able to recapture drugs identified in the discovery datasets except for VR_C3 and 4.

Conserved cell-type composition of VRS

Comparing the cell-type composition between VRS in the target dataset (BeatAML2 and TCGA), consistent with BeatAML1 (**Figure 2D**) VR_C1 showed an enrichment for more differentiated (Mono and cDC-like blasts) blasts type and depletion of primitive blast types (Prog and HSC-like blasts; **Figure S3I-J**). Enrichment for GMP-like blasts in VR_C2 relative to VR_C1 was picked up in both datasets (adjusted p-value < 0.1; **Figure S3I-J**). Enrichment for Prog-like blasts in VR_C4 was captured in both target datasets but enrichment for cDC-like blasts was not (**Figure S3I-J**). High fraction of HSC-like blasts in VR_C3 was also captured in both datasets but not an enrichment for GMP-like blasts (**Figure S3I-J**). Though there were some discrepancies, most of the strongest cluster specific cell-type patterns were conserved in both projections.

<u>Critical transcriptional characteristics are conserved in projected VRS across target datasets</u>

Analysis of differentially active pathways in VR_Cs relative to VSC and other VR_Cs also recapitulated transcriptional characteristics of VR_Cs identified in BeatAML1 (**Figure S4A-B**). in both BeatAML2 and TCGA (**Figure S4A-B**, left) VR_Cs showed a general trend of high expression of immune associated pathways relative to VSC. VR_C1 over-expression of OxPhos was captured across datasets, however the broader metabolic fitness and mTOR activation was picked up only in BeatAML2. MTORC1 signaling showed a trend towards higher activity in TCGA but did not reach significance (**Figure S4A-B**, left). Higher estrogen signaling, EMT, cell surface components we recaptured in both BeatAML2 and TCGA (**Figure S4A-B**, left). Though we observed trends towards higher angiogenesis and hypoxia pathways in VR_C3 relative to VSC, they reached significance only in BeatAML2 (**Figure S4A-B**, left). VR_C4 lacked characteristic activation of several pathways observed in other VRS relative to VSC (**Figure S4A-B**, left) as observed in BeatAML1 (**Figure 2E**), some of these pathways were even suppressed in TCGA (**Figure S4B**, left).

Comparing VR_Cs to other VR_Cs we found that VR_C1 showed consistent activation of metabolic pathways while VR_C1 and 3 showed stronger activation of inflammatory pathways (**Figure S4A-B**, right). VR_C3 specific (estrogen signaling, hypoxia, cell surface components, EMT and DNA damage but not TP53 pathway) showed higher expression in both datasets, however their specificity was more strongly preserved in TCGA, while in BeatAML2 they also showed trends for higher expression in VR_C1 relative to VR_C2 and 4(**Figure S4A-B**, right). Interestingly, though VR_C1 and 3 showed high expression of inflammatory pathways, as in beatAML1 (**SuppFig3 K-L**), VR_C3 showed much higher degree of CTL (**Figure S4C)** infiltration and higher expression of cytotoxic effector and checkpoint genes (**Figure S4D**). We also recaptured lower expression of HOX genes in VR_C2 relative to other VRS, except in the case of VSC in TCGA (**Figure S4D**).
